# A non-retinoid triazolopyrimidine RBP4 antagonist for the treatment of Stargardt disease

**DOI:** 10.64898/2026.09.14.751502

**Authors:** K. Alison Rinderspacher, Andras Varadi, Boglarka Racz, Andrew S. Wasmuth, Shi-Xian Deng, Patricia Weber, Donald W. Landry, Peter R. Bernstein, Konstantin Petrukhin

## Abstract

Stargardt disease is a juvenile-onset retinal dystrophy characterized by the buildup of cytotoxic lipofuscin deposits in the retinal pigment epithelium (RPE), leading to photoreceptor degeneration and eventual blindness. Currently, there are no FDA-approved treatments for Stargardt disease. Bisretinoids, byproducts of the visual cycle, are the major cytotoxic components of the lipofuscin deposits, and bisretinoid synthesis relies on the traffic of retinol from the bloodstream to the retina. Selective targeting of the key retinol transporter, Retinol-Binding Protein 4 (RBP4), offers an appealing strategy for halting the buildup of lipofuscin in the RPE and arresting the progression of Stargardt disease. Retinol delivery depends on RBP4 interaction with another serum protein, Transthyretin (TTR). We previously reported several libraries of RBP4 antagonists that effectively blocked the association of the TTR-RBP4-retinol tertiary complex, thereby lowering the overall retinol load in the retina; however, some chemotypes displayed off-target activity that warranted further optimization. Here, we report the pharmacological characterization of AKR-XI-85 and its analogs as promising non-retinoid small-molecule RBP4 antagonists. AKR-XI-85 displayed excellent *in vitro* and *in vivo* efficacy and desirable pharmacokinetic properties without any limiting off-target activity. In *Abca4^-/-^* mice, chronic dosing of the compound induced a prolonged reduction in serum RBP4 levels and achieved a dramatic, 70 % reduction in the accumulation of A2E, a critical component of toxic lipofuscin. As such, AKR-XI-85 may be an attractive drug candidate for the treatment of Stargardt disease and other lipofuscin-dependent retinopathies.

## 1. Introduction

Stargardt disease (STGD1; MIM 248200) is the most common juvenile-onset macular dystrophy with an estimated prevalence of 1:8,000-10,000 in the United States.(1, 2) STGD1 is characterized by bilateral macular atrophy that leads to progressive loss of retinal function and subsequent decrease in central visual acuity. Another characteristic of the disease is the presence of yellow-white flecks at the posterior pole at the level of the retinal pigment epithelium (RPE).(3, 4) While most patients develop STGD during childhood or early into adulthood, the age of onset is highly variable but a later age of onset is typically associated with a better prognosis.(5–7) STGD1 is caused by the mutations of a single gene, ABCA4, and shows autosomal recessive inheritance.(8, 9) ABCA4 belongs to the family of ATP-binding cassette transporters expressed in photoreceptors. Its primary role is to facilitate the removal of *N*-retinylidene-phosphatidylethanolamine from disk membranes of rod and cone photoreceptor cells.(10) This transport activity prevents the buildup of toxic retinaldehydes and their bisretinoid condensation products. Defective ABCA4 in STGD1 leads to the accumulation of cytotoxic lipofuscin comprising bisretinoids, such as the pyridinium compound A2E in RPE cells.(11) Currently, there are no FDA-approved treatments for STGD1.(12) Developing a pharmacological treatment capable of inhibiting the buildup of lipofuscin and subsequently preventing or halting the progression of retinal atrophy would address a critical unmet medical need. The visual cycle is fueled by retinoids, which are constantly supplied to the retina from the circulation. Retinoids are found at high levels in the body, especially in the liver.(13) Retinoids are transported from hepatic stores to other organs mainly as all-*trans*-retinol bound to Retinol-Binding Protein 4 (RBP4), a low-molecular-weight (21 kDa) lipocalin protein.(14) In the serum, retinol-bound RBP4 is found as a tertiary complex with transthyretin (TTR, prealbumin), which stabilizes the retinol-RBP4 complex and prevents its renal filtration.(15) Retinol binding is required for the formation of the TTR-RBP4 complex; apo-RBP4 (retinol-free RBP4) has a low affinity for TTR and is rapidly removed from the circulation through glomerular filtration and subsequent catabolism.(16) Therefore, the disruption of the RBP4-TTR interaction presents an attractive approach to limiting the retinol load of the retina, thereby reducing the levels of potential A2E precursors. Fenretinide, a synthetic analog of all-*trans*-retinol (Fig. 1) competes with retinol to bind RBP4 and is therefore able to block the formation of the RBP4-TTR complex.(17, 18) Amgen introduced the first non-retinoid RBP4 antagonist, A1120 (Fig. 1), which was tested for its potential as a diabetes therapy but did not show efficacy in diabetes models.(19) In pursuit of a treatment for macular degeneration, we have previously developed and characterized several classes of non-retinoid and selective RBP4 antagonists.(20–25) We showed that our selective RBP4 antagonists induce desirable decreases in serum RBP4 in several species.(21–25) In particular, BPN-14136, a heterocyclic carboxylic acid with a [3.3.0]-octahydrocyclopenta[*c*]pyrrolo bicyclic core (Fig. 1), displayed outstanding RBP4 reduction, robust inhibition of bisretinoid formation in a murine model of lipofuscinogenesis, and had desirable pharmacokinetic (PK) and pharmacodynamic (PD) properties in mice, rats, dogs and non-human primates.(22, 23, 25) BPN-14136 was also able to normalize complement system dysregulation, an important contributor to photoreceptor degradation in macular degeneration.(23) However, the carboxylic acid featured in the BPN-14136 series may represent a potential liability. Certain drugs containing carboxylic acid have been linked to idiosyncratic drug toxicity, possibly due to the formation of reactive acyl glucuronide metabolites.(26–28) Furthermore, the aromatic carboxylic acid group featured in BPN-14136 is likely to be involved in mediating the appreciable off-target activity of this compound as a PPARγ agonist. This PPARγ agonistic activity can significantly complicate the development of BPN-14136 as a drug candidate for STGD, given that PPARγ activation is associated with increased risk of death, myocardial infarction, stroke, congestive heart failure, hepatotoxicity, peripheral edema, weight gain, and carcinogenicity.(29–32) In this study, we report the optimization of BPN-14136 with the primary goal of preserving or improving the *in vitro* potency of BPN-14136 while eliminating the off-target PPARγ agonistic activity arising from the presence of the aromatic carboxylic acid group. Our efforts have led to the identification of AKR-XI-85 (Fig. 1), a 1,2,4-triazolopyrimidinyl amide analog of BPN-14136, which showed excellent *in vitro* and *in vivo* potency and optimal drug-like properties.

**Figure 1.**
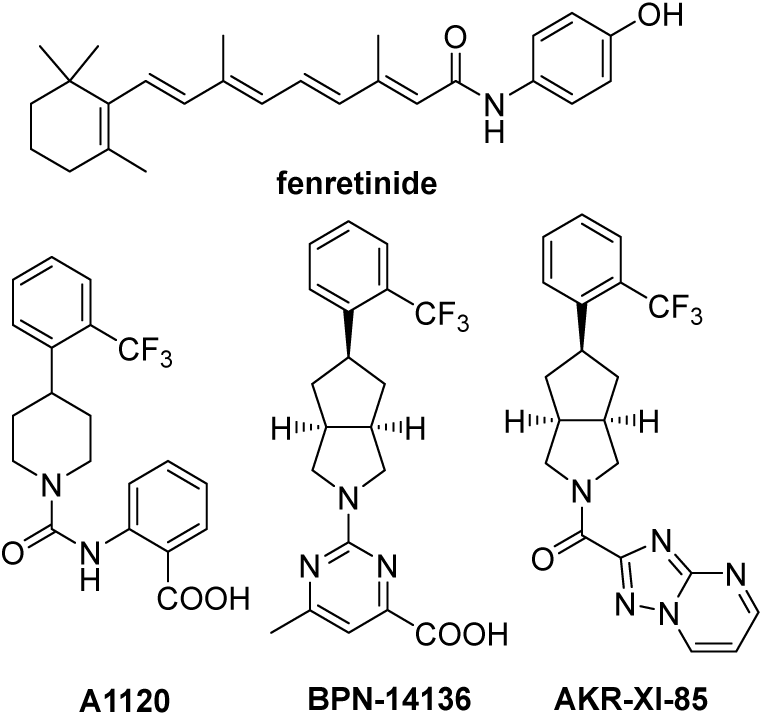
Chemical structure of RBP4 ligands fenretinide, A1120, BPN-14136, and AKR-XI-85.

## 2. Materials and Methods

### In vitro SAR assays

#### *In vitro* RBP4 binding assay

RBP4 binding was determined using a scintillation proximity assay (SPA) as described previously.(33) The assay quantified displacement of [³H]-all-*trans*-retinol from human RBP4 (urine-derived; Fitzgerald, 30R-AR022L). RBP4 was biotinylated with the EZ-link Sulfo-NHS-LC-Biotin kit (ThermoFisher, #21335). Reactions (100 μL) were carried out in SPA buffer (1× PBR, pH 7.4, 1 mM EDTA, 0.1% BSA, 0.5% CHAPS) containing 10-nM [³H]-retinol (48.7 Ci/mmol; PerkinElmer), 0.3 mg/well Streptavidin-PVT beads (PerkinElmer, RPNQ0006), and 50-nM biotinylated RBP4. Nonspecific binding was assessed with 20-μM unlabeled retinol (Sigma, #95144). Plates were incubated for 16 h at room temperature with gentle agitation, and radioactivity was recorded on a CHAMELEON plate reader (Hidex, Turku, Finland).

#### RBP4-TTR interaction assay

Antagonist activity of analogs toward the all-*trans*-retinol–dependent RBP4–TTR interaction was assessed using a homogeneous time-resolved fluorescence (HTRF) assay as we described. previously.(33) Untagged TTR (Calbiochem, #529577) and maltose-binding protein (MBP)–tagged RBP4 expressed in *E. coli* were employed. TTR was labeled with Eu³⁺ Cryptate using the HTRF Cryptate kit (Cisbio, #62EUSPEA). Reactions (16 μL) were prepared in buffer containing 10 mM Tris-HCl (pH 7.5), 1 mM DTT, 0.05% NP-40, 0.05% Prionex, 6% glycerol, and 400 mM KF, with 60 nM MBP-RBP4, 5 nM TTR-Eu, 26.7 nM anti-MBP antibody–d2 conjugate (Cisbio, #61MBPDAA), and 1 μM all-*trans*-retinol (Sigma, #95144). Reactions were carried out under dim red light and incubated overnight at 4 °C. Fluorescence was read on a SpectraMax M5e (Molecular Devices) at 337 nm excitation, with emissions at 668 and 620 nm (75 μs delay). The HTRF signal was expressed as Flu668/Flu620 × 10,000.

#### Assay for PPARγ agonism

For bacterial expression, the ligand-binding domain of PPARγ (amino acids 176-477, GenBank accession number NP_005028) was subcloned into the SalI-NotI sites of pGEX-6p–3. After introduction of expression plasmids to the BL21-Gold(DE3)pLysS E.coli strain (Stratagene), GST-tagged PPARγ-LBD was purified from 1-L cultures using an AKTA FPLC system (GE Healthcare) equipped with 5-mL GST Trap HP. Peptide N-COR-2, Biotin-(aminohexanoic acid)-ADPASNLGLEDIIRKALMGSF-NH2, representing the second nuclear receptor interacting fragment of corepressor N-COR was synthesized by Genemed Synthesis, Inc. The assay was performed in white Costar 384-well polystyrene plates (Corning, Corning, NY) in 16 μL final volume. The final composition of 1X HTRF buffer (Assay Buffer) was 10 mM Tris-HCl, pH 7.5, 100 mM potassium fluoride (KF), 0.05% *w/v* bovine serum albumin (BSA), 0.05% NP-40, 1 mM dithiothreitol (DTT), 6% Glycerol. 10 μL of a mix containing GST-tagged PPARγ-LBD and biotinylated N-COR-2 peptide in 1X HTRF buffer were added to 4 μL of a compound dilution in 1X buffer. After the 16-hour incubation at 4°C, 2 μL of the detection mix were added to a well. Final concentrations were 7.0 nM GST-PPARγ, 300 nM Biotin-N-COR-2, 0.75 nM Europium cryptate-anti-GST-antibody (Eu(K)-anti-GST Ab (CisBio, Bedford, MA) and 42 nM Streptavidin-XL665 (SA-XL665; CisBio, Bedford, MA). Plates were incubated at 4°C for an additional 16-24 hours followed by HTRF measurement on SpectraMax M5e Multimode Plate reader (Molecular Devices, Sunnyvale, CA). Two readings were taken: Reading 1 for time-gated energy from Eu(K) to XL665 (337 nm excitation, 668 emission, counting delay 50 µsec, counting window 400 µsec) and Reading 2 for Eu(K) time-gated fluorescence (337 nm excitation, 620 nm emission, counting delay 50 µsec, counting window 400 µsec). The signal was expressed as the ratio of fluorescence intensity according to Equation 1: 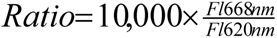 where Fl668nm represents the measured fluorescence emission at 668nm and Fl620nm represents the measured fluorescence emission at 620nm.

### *In vitro* ADME tests

*In vitro* ADME tests were conducted at Absorption Systems and Eurofins Discovery. Plasma protein binding for AKR-XI-85 was determined (in triplicates) by equilibrium dialysis of plasma against phosphate buffered saline (pH 7.4). Plasma spiked with AKR-XI-85 at a concentration of 1 μM was loaded to one side (donor) of the dialysis device insert, and phosphate buffered saline was loaded to the other side (receiver). After four hours, the concentration of AKR-XI-85 was assessed in both the donor and receiver sides.

Metabolic stability determinations for AKR-XI-85 and testosterone (positive control) were conducted in the presence of human, dog, and cynomolgus monkey liver microsomes. All measurements were done in duplicate. Pooled mixed gender human donor microsomes, male beagle dog microsomes and male cynomolgus monkey were obtained from BioIVT (Baltimore, MD). AKR-XI-85 was prepared as a 10 mM stock solution in DMSO. A mixture containing 50 mM potassium phosphate buffer pH 7.4 and 1 mg/mL liver microsomes was pre-warmed for 10 min at 37°C in a shaking water bath, followed by the addition of test compound. Final DMSO concentration in the reaction mix was 0.1%. Reactions with cofactor were initiated by adding an NADPH-regenerating system to the incubation mixtures (final concentrations of 1.3 mM NADP^+^, 3.3 mM glucose-6-phosphate, and 0.4 U/mL glucose-6-phosphate dehydrogenase). The final volume of the reaction mixture was 800 μL, containing 1 mg/mL liver microsomes, and 1 μM test compound. Aliquots (100 μL) of reaction mixtures were removed from the incubation plate at pre-defined timepoints and mixed with 150 μL of ice-cold acetonitrile, incubated on ice for 15 min, and samples were centrifuged (3,600 rpm, 10 min, 4°C) to precipitate protein. The supernatants were diluted 1:1 (v/v) with water containing the internal standard and subjected to LC/MS analysis. The low limit of quantitation for AKR-XI-85 was 0.01 µg/ml.

Kinetic aqueous solubility determination in PBS (pH 7.4) was conducted using UV detection (230 nm). Aqueous solubility (μM) was determined by comparing the peak area of the principal peak in a calibration standard (200 μM) containing organic solvent (methanol/water, 60/40, v/v) with the peak area of the corresponding peak in a buffer sample. In addition, chromatographic purity (%) was defined as the peak area of the principal peak relative to the total integrated peak area in the HPLC chromatogram of the calibration standard.

### Animal experiments

Animal studies were approved by the Institutional Animal Care and Use Committee (IACUC) of Columbia University and performed following the guidelines of the Association for Research in Vision and Ophthalmology (ARVO) statement for the “*Use of Animals in Ophthalmic and Vision Research”*.

### Pharmacokinetic and pharmacodynamic studies

Pharmacokinetic studies were conducted in CD-1 mice and Beagle dogs at Absorption Systems. Mice and dogs received a single IV dose of a test compound at 2 mg/kg or a single PO dose of a test compound at 5 mg/kg. IV vehicle was 3% DMA/45% PEG300/12% ethanol/40% sterile water; PO vehicle was 2% Tween 80 in 0.9% saline. Blood was collected from mice and dogs at pre-dose and at 8–9 timepoints over 48 hours following compound administration. Following protein extraction with acetonitrile, compound levels in whole blood were measured by LC-MS/MS. The mean plasma drug level values were analyzed using a WinNonlin package by noncompartmental modeling with the sparse sampling feature.

Albino Balb/c mice were used for the pharmacodynamic study. Mice received single escalating doses of AKR-XI-85 to evaluate the effects of the compound on serum RBP4 dynamics. Blood samples were collected at pre-dose and at six timepoints over 24 hours following oral gavage administration of AKR-XI-85 at 15, 25, or 35 mg/kg. The compound was formulated in 2 % Tween 80 (v/v) prepared in 0.9% saline. Whole blood (∼10 μL) was collected into centrifuge tubes, allowed to clot at room temperature for 30 min, and centrifuged at 2000 min for 15 min at 4 °C to obtain serum for RBP4 measurements.

### Mouse serum RBP4 measurements

In the *Abca4^−/−^* dosing experiment, blood samples were collected from a tail vein at pre-dose and after 60 days of AKR-XI-85 administration. In mouse PD experiments, blood was collected as described above. Following blood clotting and serum preparation, serum RBP4 was measured using the RBP4 (mouse/rat) dual ELISA kit (AdipoGen, Switzerland) following the manufacturer’s instructions.

### Assessment of bisretinoid-lowering efficacy in the *Abca4^-/-^* mouse model: AKR-XI-85 dosing, bisretinoid extraction and analysis

*Abca4^−/−^* mice (129/SV × C57BL/6J), homozygous for the Rpe65-Leu450, were bred as described previously(34, 35) and used to assess the AKR-XI-85 effect on bisretinoid accumulation. AKR-XI-85 was formulated into Purina 5035 rodent chow at Research Diets, Inc. (New Brunswick, NJ) to ensure consistent 35 mg/kg daily oral dosing over a period of 60 days. Following the dosing regimen, posterior eyecups were prepared from mouse eyes. Lipofuscin bisretinoids were isolated from posterior eyecups of untreated wild-type mice (9 eyes), vehicle-treated *Abca4^−/−^*mice (12 eyes), and AKR-XI-85–treated *Abca4^−/−^* mice (12 eyes). Eyecups were pooled and homogenized in PBS with a tissue grinder. Chloroform/methanol (2:1, v/v) was added in an equal volume, and the samples were extracted three times. The combined organic phases were evaporated under a gentle stream of argon and reconstituted in 100 μL of methanol. A2E was separated on a C18 reverse-phase column using a methanol–water gradient (85–96% methanol containing 0.1% trifluoroacetic acid). Absorbance at 430 nm was monitored with a photodiode array (PDA) detector, and the A2E levels were quantified from peak areas using a calibration curve generated with synthetic A2E standards.

### Metabolite Identification of AKR-XI-85

Metabolite Identification studies were performed by Absorption Systems, employing the following conditions: Mixed-gender human liver microsomes (Lot# 1710084) were purchased from XenoTech. The reaction mixture, minus NADPH, was prepared as described below. The test article (TA) was added into the reaction mixture at a final concentration of 50 μM. An aliquot of the reaction mixture (without cofactor) was equilibrated in a shaking water bath at 37 °C for 3 minutes. The reaction was initiated by the addition of the cofactor, and the mixture was incubated in a shaking water bath at 37 °C. Aliquots (100 μL) were withdrawn at 0 and 60 minutes. TA samples were immediately combined with 400 μL of ice-cold 50/50 MeCN/H_2_O containing 0.1 % formic acid and an internal standard to terminate the reaction. The samples were then mixed and centrifuged to precipitate proteins. A Dionex XR3000 quaternary solvent HPLC system, equipped with a column compartment thermostat, set to 40 °C, was used for chromatographic separation (Column: BEH C18 XBridge, 2.5 μm, 100 × 2.1 mm (Waters)). A linear gradient from 5 % to 100 % B over 45 min, then 100 % B for 0.1 min (A = 0.1 % acetic acid + H_2_O, B = CH_3_CN)) was employed.

The Equilibrium Solubility, hERG, PXR Activation, MDCKII-MDR1 Permeability, Caco-2 Permeability, and CYP Phenotyping assays were performed by Absorption Systems and Eurofins Discovery following their standard protocols.

### Synthesis of AKR-XI-85

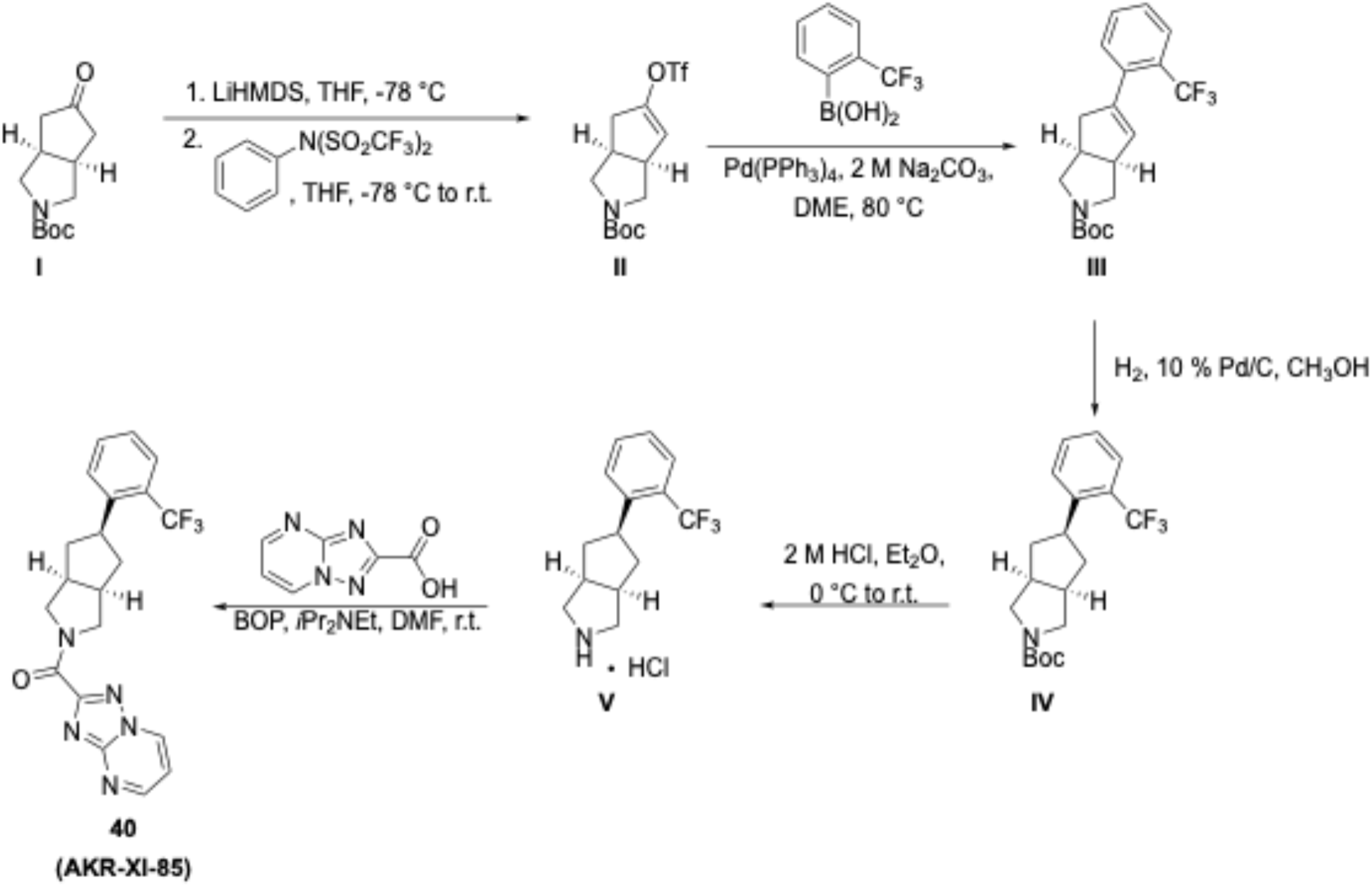

#### Experimental

Melting points were determined on a Mel-Temp II Laboratory Devices apparatus and are reported uncorrected. ^1^H NMR and ^13^C NMR spectra were recorded on an Agilent 400-MR 400-MHz NMR spectrometer, operating at 400 MHz (^1^H NMR) and 101 MHz (^13^C NMR). Chemical shifts are reported in parts per million using the residual proton or carbon signal (CDCl_3_: δ_H_ 7.26, δ_c_ 77.16) as an internal reference. The apparent multiplicity (s = singlet, d = doublet, t = triplet, q = quartet, m = multiplet) and coupling constants (in Hz) are reported in that order in the parentheses after the chemical shift. Liquid chromatography and mass spectrometry were performed on a Shimadzu 2020 UFLC mass spectrometer, using a Waters Sunfire column (C18, 5μm, 2.1 mm x 50 mm, a linear gradient from 5 % to 100 % B over 15 min, then 100 % B for 2 min (A = 0.1 % formic acid + H_2_O, B = 0.1 % formic acid + CH_3_CN), flow rate 0.2000 mL/min). HRMS (EI) was performed by Furong Sun and associates at the University of Illinois (Urbana-Champaign) and the results were determined to be within ±0.4 % of the theoretical values. All reagents and solvents were used as received from major commercial suppliers, such as Sigma-Aldrich, Fisher Scientific, and Alfa Aesar without further purification. All air-or moisture-sensitive reactions were run under an atmosphere of argon in oven-dried glassware unless otherwise noted.

**Step 1:** Lithium bis(trimethylsilyl)amide (LiHMDS) was added via syringe to a yellow-orange solution of *tert*-butyl-5-oxooctahydrocyclopenta[*c*]pyrrole-2-carboxylate (**I**) (1.4 M) in anhydrous tetrahydrofuran (THF), stirred at -78 °C. The reaction mixture was stirred at -78 °C for 1h 45 min, after which a solution of *N*-phenyltrifluoromethanesulfonimide (0.9 M) in anhydrous THF was added portionwise. The reaction mixture was stirred at -78 °C for another 2 h, after which it was allowed to warm to room temperature. It was concentrated *in vacuo* and purified via normal phase silica gel column chromatography (0 % to 15 % ethyl acetate in hexanes).

**Step 2:** Compound **II** (0.04 M) (1 equiv) and 2-(trifluoromethyl)phenylboronic acid (2.5 equiv) were stirred in a 1:2 mixture of 2 M aqueous sodium carbonate and 1,2-dimethoxyethane. The reaction mixture was evacuated and purged with argon.

Tetrakis(triphenylphosphine)palladium(0) (0.1 equiv) was added, and the reaction mixture was evacuated and purged with argon. It was heated to and stirred at 80 °C for 6 h, after which it was allowed to cool to room temperature. Ethyl acetate was added and the reaction mixture was concentrated *in vacuo*. An additional volume of ethyl acetate was added. The organic and aqueous layers were separated. The organic layer was washed with brine (2x) and dried with anhydrous sodium sulfate. The solvent was evaporated *in vacuo*. The resulting crude material was purified via normal phase silica gel column chromatography (hexanes followed by 20 % ethyl acetate in hexanes followed by ethyl acetate).

**Step 3:** Compound **III** (0.5 M) was stirred in methanol. The reaction mixture was evacuated and purged with argon using a dual manifold. 10 % Palladium on carbon was added. The reaction mixture was evacuated and purged with argon. Then it was evacuated and purged three times with hydrogen, using a balloon, fitted with a three-way adapter, after which a steady stream of hydrogen was allowed to pass through the flask. The reaction mixture was stirred overnight, then filtered through a Celite pad with methanol. The filtrate was concentrated *in vacuo*. The resulting crude material was carried on to the next step.

**Step 4:** Compound **IV** (0.7 M) (1 equiv) was stirred in methylene chloride at 0 °C. A 2-M solution of HCl in diethyl ether (5.6 equiv) was added portionwise. The reaction mixture was allowed to warm to room temperature and was stirred overnight. Subsequent addition of diethyl ether resulted in the formation of a white precipitate, which was isolated via vacuum filtration.

**Step 5:** The hydrochloric acid salt, **V**, (0.14 M) (1 equiv), [1,2,4]triazolo[1,5-*a*]pyrimidine-2-carboxylic acid (1 equiv), and (benzotriazol-1-yloxy)tris(dimethylamino)phosphonium hexafluorophosphate (BOP) (1.5 equiv) were stirred in anhydrous DMF at room temperature. Diisopropylethylamine (3.0 equiv) was added via syringe. The reaction mixture was stirred overnight, after which distilled water was added. The resulting off-white precipitate was isolated via vacuum filtration.

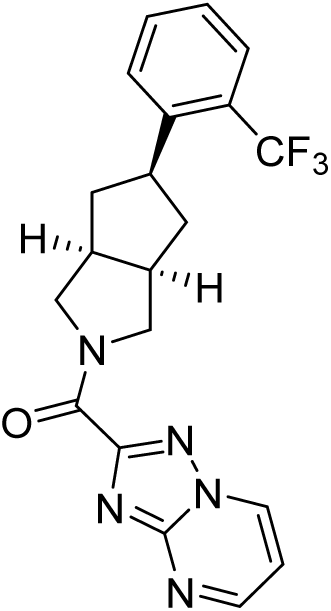

**AKR-XI-85** (**40**): Off-white solid; Yield: 87 %; 1H NMR (400 MHz, (CDCl3)): δ 8.95 (d, *J* = 6.8 Hz, 1H), 8.92 (dd, *J* = 3.6, 1.6 Hz, 1H), 7.60 (d, *J* = 8.0 Hz, 1H), 7.51 (d, *J* = 6.0 Hz, 2H), 7.28 (d, *J* = 7.2 Hz, 1H), 7.24 (dd, *J* = 6.8, 4.8 Hz, 1H), 4.17 (s, 2H), 3.93 (d, *J* = 5.6 Hz, 2H), 3.53 (septet, *J* = 6.8 Hz, 1H), 2.91 (sextet, *J* = 6.4 Hz, 2H), 2.37 (doublet of pentets, *J* = 37.6, 6.8 Hz, 2H), 1.69-1.57 (m, 2H); 13C NMR (101 MHz, (CDCl3)): δ 161.6, 159.4, 155.7, 155.0, 142.9, 136.5, 132.3, 128.0 (2C), 126.2, 125.8 (2C), 111.5, 54.2, 52.6, 44.1, 43.0, 41.6, 41.4, 41.1; LC-MS (M^+^+H): 402; EI+ HRMS (*m*/*z*): [M]+ calcd. for C20H18N5OF3: 401.14635, Found: 401.14615.

#### Molecular modeling

Docking and conformation analyses were conducted in the Molecular Operating Environment (MOE) as implemented by Chemical 3Computing Group (CCG), Montreal Canada. RBP4 structures were taken from the Protein Data Bank (PDB).(36)

## 3. Results

### 3.1 BPN-14136 acts as a PPARγ agonist in the *in vitro* co-repressor release assay

As we previously reported, BPN-14136 demonstrates potent RBP4 binding (IC_50_=12.8 nM) and effectively disrupts the RBP4 interaction with transthyretin (IC_50_=43.6 nM). (22) BPN-14136 lacks inhibition of standard CYP 450 enzymes, shows good stability in liver microsomes (22), displays favorable pharmacokinetic properties (23, 25), and successfully improves the phenotype in a Stargardt disease mouse model.(23) While BPN-14136 shows no off-target activity at the hERG channel or within a standard CEREP selectivity screening panel of fifty-five GPCRs, enzymes, ion channels, and transporters (22), we opted to further assess its selectivity in an *in vitro* assay for detecting PPARγ agonistic activity. The PPARγ ligand-binding domain consistently interacts with transcriptional corepressors N-COR and SMRT in the absence of an agonistic ligand (37), while binding of an agonist leads to the release of the transcriptional co-repressor. We previously developed an *in vitro* HTRF assay to measure the agonist-induced release of the biotinylated N-COR-2 peptide from the purified PPARγ fragment. (38) Dose titration of BPN-14136 in the PPARγ activity assay revealed its significant agonistic activity (IC_50_=1.9 µM; Fig. 2). The control PPARγ agonist rosiglitazone was significantly more potent (IC_50_=83 nM). As shown in Fig. 3, altering the carboxylic acid group in the BPN-14136 analogs leads to the elimination or significant decrease in PPARγ agonistic activity. This suggests that this part of the molecule plays a crucial role in modulating PPARγ agonistic activity. Interestingly, the *exo*-isomer of BPN-14136 (compound BPN-14815) exhibits markedly reduced RBP4 binding activity, yet it demonstrates equivalent potency to the *endo*-isomer BPN-14136 in the PPARγ assay. (Fig. 3). It seemed reasonable to suggest that designing BPN-14136 analogs without a carboxylic acid group could result in compounds without undesired PPARγ agonistic effects.

**Figure 2.**
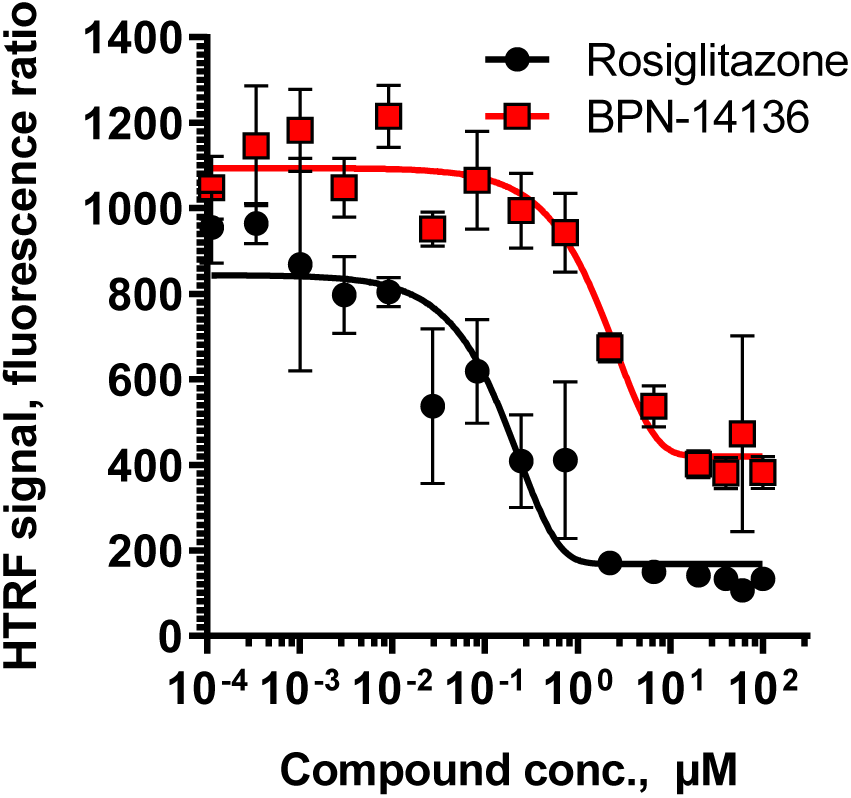
Characterization of BPN-14136 in the PPARγ-N-CoR interaction assay. BPN-14136 dose-dependently decreases the interaction between the N-CoR co-repressor fragment and PPARγ ligand-binding domain, confirming the PPARγ agonist activity of the test compound. Rosiglitazone is used as a positive control. Data represented as the mean ±SD with three independent dose titration experiments performed. HTRF signal is expressed as fluorescence ratio (Fl_668_/Fl620 x 10,000).

**Figure 3.**
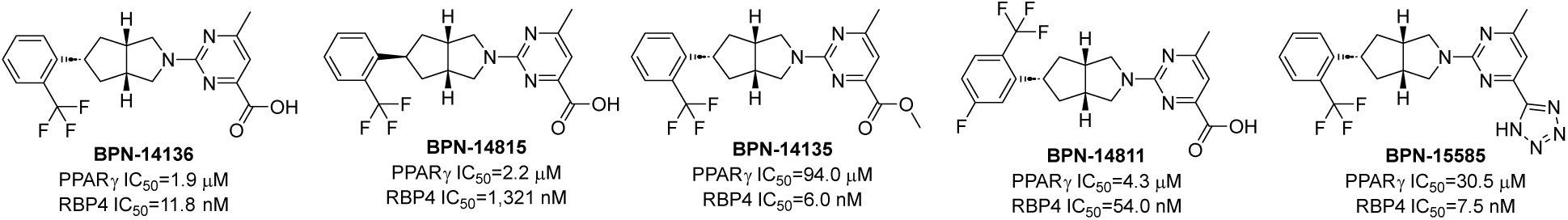
PPARγ agonistic activity and RBP4 SPA binding affinity for BPN-14136 analogs. The aryl carboxylic group present in BPN-14136 and BPN-14815 but absent in BPN-15585 and BPN-14135 seems to be involved in modulating the activity of BPN-14136 as a PPARγ agonist

### 3.2 Medicinal chemistry optimization and identification of AKR-XI-85

Our primary objective was to create a series of analogs lacking the carboxylic acid group on the lower aromatic ring of BPN-14136, while ensuring that these compounds maintain sufficient potency in *in vitro* RBP4 binding and RBP4-TTR interaction assays. Initially, we synthesized ester and amide derivatives of BPN-14136 that retained the pyrimidine bottom group of the parent molecule (Table 1). *In vitro* RBP4 binding potency was tested in the SPA assay and the compounds’ ability to block the retinol-dependent TTR-RBP4 interaction was assayed in the TTR-RBP4 HTRF assay. (21, 22) Esterification of the carboxylic acid group bound to the pyrimidine bottom ring affected the *in vitro* RBP4 binding potency of the compounds negatively. While analogs bearing smaller alkyl groups such as an isopropyl (**1**, IC_50_ = 15 nM), 4-hydroxybutyl (**9**, IC_50_ = 33 nM), a 3-hydroxypropyl (**10**, IC_50_ = 22 nM), a methylhydroxyacetyl (**17**, IC_50_ = 17 nM), and a methoxyethyl (**18**, IC_50_ = 19 nM) group retained good RBP4 binding potency in the SPA assay, bulkier alkyl groups reduced the binding affinity significantly. This effect is most notable in the case of the *tert*-butyl (**11**, IC_50_ = 318 nM) and 4-methoxybenzyl (**15**, IC_50_ = 112 nM) esters. A similar trend could be observed in the derivatives’ ability to block the retinol-dependent TTR-RBP4 interaction. Esterification, however, practically eliminated the potency of all compounds with the isopropyl ester (**1**) being the best in this series with a meager IC_50_ of 210 nM. Changing the methyl group to an electron-withdrawing chlorine on the pyrimidine ring (**4, 6, 7**) reduced the RBP4 binding potency. Amides (**2, 3, 16, 25, 26**) displayed moderate binding affinities with poor potency in the TTR-RBP4 HTRF assay. Replacing the carboxylic acid group with an amino moiety along with its acylated version diminished the *in vitro* potency.

**Table 1.**
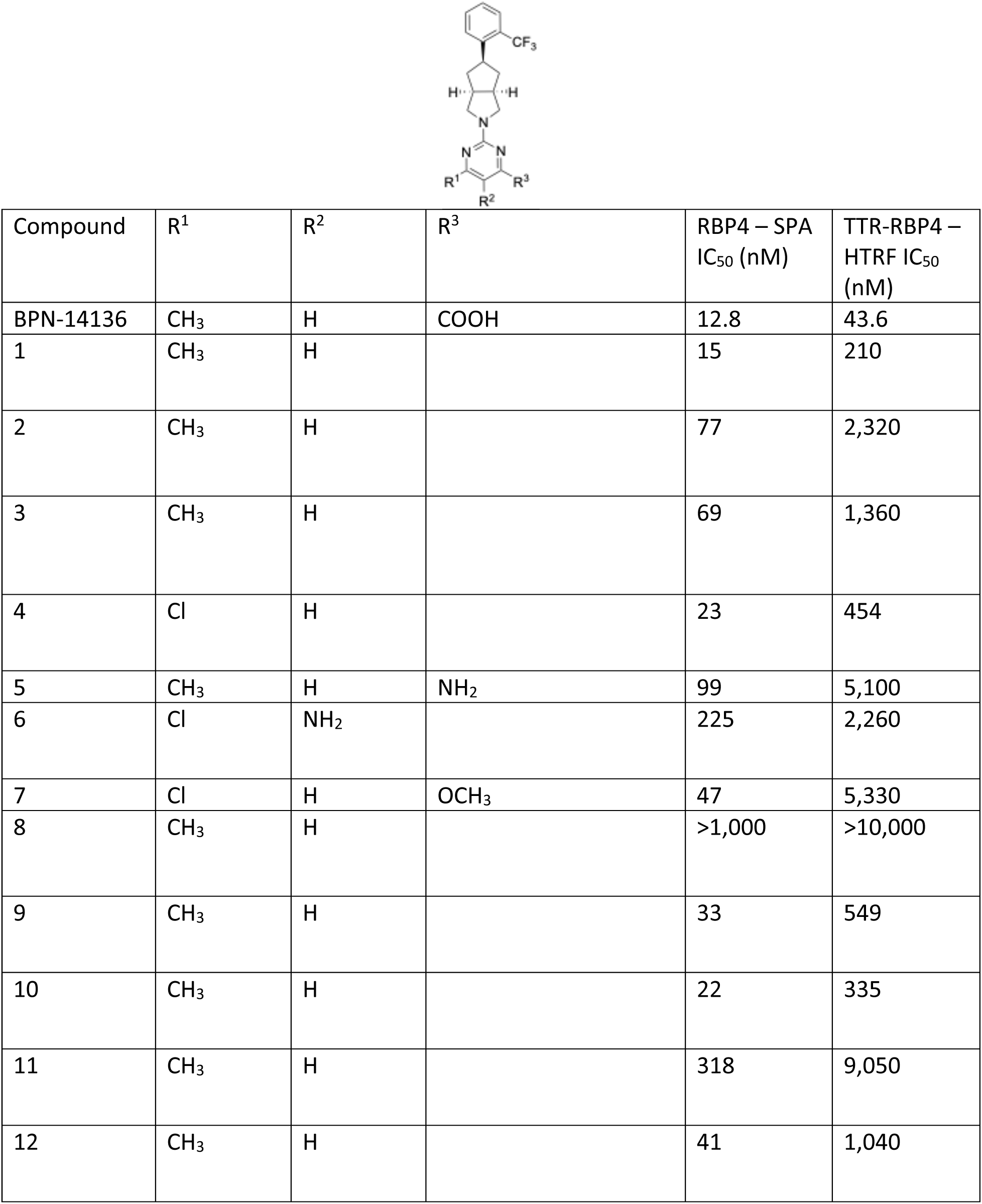
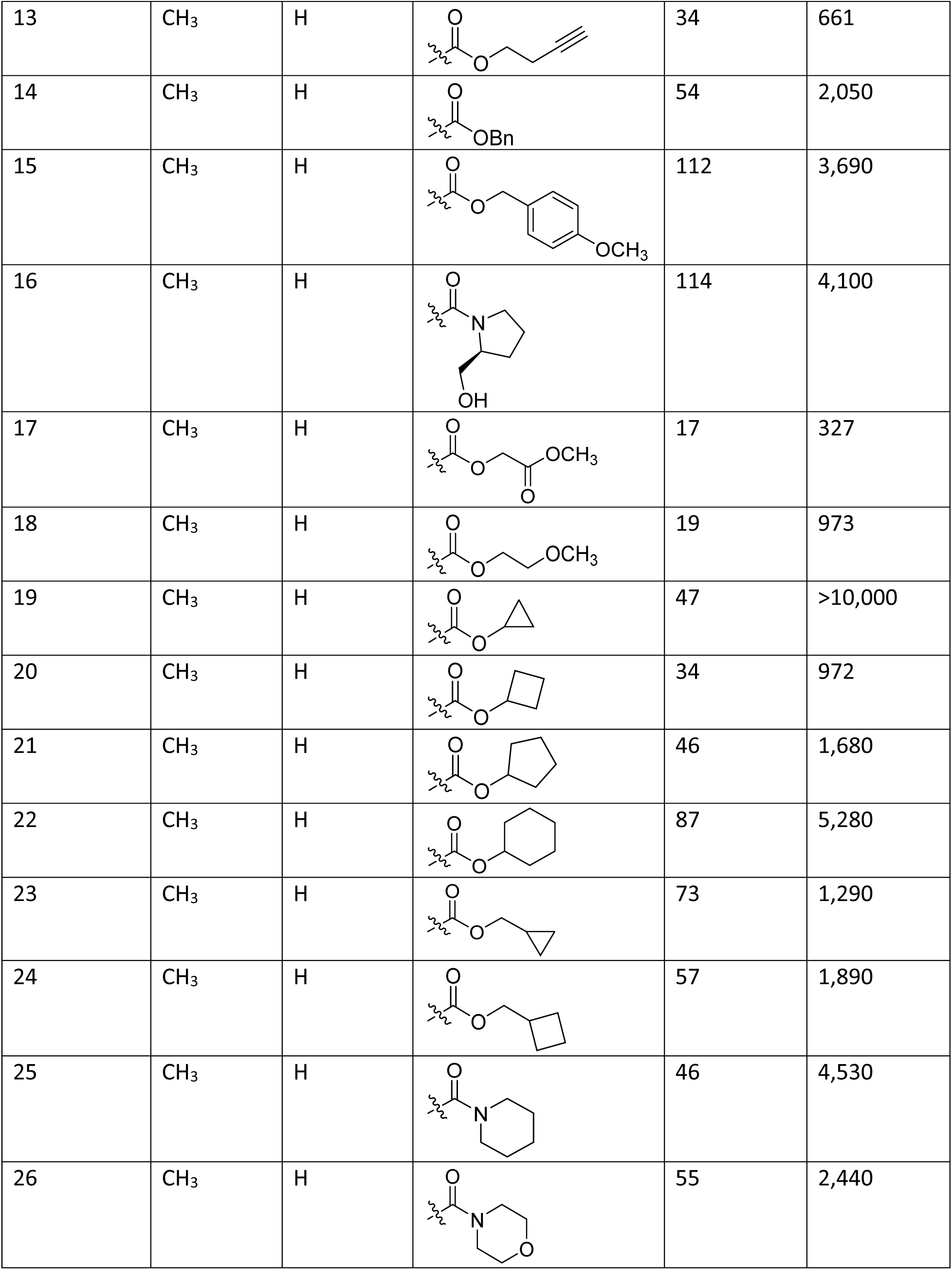
Substituted pyrimidine analogs of BPN-14136.

| Compound | R<sup>1</sup> | R<sup>2</sup> | R<sup>3</sup> | RBP4 – SPA<br>IC<sub>50</sub> (nM) | TTR-RBP4 –<br>HTRF IC<sub>50</sub><br>(nM) |
| --- | --- | --- | --- | --- | --- |
| BPN-14136 | CH<sub>3</sub> | H | COOH | 12.8 | 43.6 |
| 1 | CH<sub>3</sub> | H |  | 15 | 210 |
| 2 | CH<sub>3</sub> | H |  | 77 | 2,320 |
| 3 | CH<sub>3</sub> | H |  | 69 | 1,360 |
| 4 | Cl | H |  | 23 | 454 |
| 5 | CH<sub>3</sub> | H | NH<sub>2</sub> | 99 | 5,100 |
| 6 | Cl | NH<sub>2</sub> |  | 225 | 2,260 |
| 7 | Cl | H | OCH<sub>3</sub> | 47 | 5,330 |
| 8 | CH<sub>3</sub> | H |  | >1,000 | >10,000 |
| 9 | CH<sub>3</sub> | H |  | 33 | 549 |
| 10 | CH<sub>3</sub> | H |  | 22 | 335 |
| 11 | CH<sub>3</sub> | H |  | 318 | 9,050 |
| 12 | CH<sub>3</sub> | H |  | 41 | 1,040 |
| 13 | CH<sub>3</sub> | H |  | 34 | 661 |
| 14 | CH<sub>3</sub> | H |  | 54 | 2,050 |
| 15 | CH<sub>3</sub> | H |  | 112 | 3,690 |
| 16 | CH<sub>3</sub> | H |  | 114 | 4,100 |
| 17 | CH<sub>3</sub> | H |  | 17 | 327 |
| 18 | CH<sub>3</sub> | H |  | 19 | 973 |
| 19 | CH<sub>3</sub> | H |  | 47 | >10,000 |
| 20 | CH<sub>3</sub> | H |  | 34 | 972 |
| 21 | CH<sub>3</sub> | H |  | 46 | 1,680 |
| 22 | CH<sub>3</sub> | H |  | 87 | 5,280 |
| 23 | CH<sub>3</sub> | H |  | 73 | 1,290 |
| 24 | CH<sub>3</sub> | H |  | 57 | 1,890 |
| 25 | CH<sub>3</sub> | H |  | 46 | 4,530 |
| 26 | CH<sub>3</sub> | H |  | 55 | 2,440 |

Next, in an attempt to develop novel chemical matter with improved potency, we created a series of compounds in which the substituted bottom pyrimidine ring was replaced with various heteroaromatic groups (Table 2). Compounds exhibiting high potency in the RBP4 binding assay were evaluated in the PPARγ assay. The tested compounds **30, 32, 40, 43, 44** did not display any affinity (IC_50_ > 100 µM) for PPARγ (data not shown). The triazolopyrimidine analog **40** (AKR-XI-85) displayed very good RBP4 binding potency (IC_50_ = 23 nM) in the SPA assay and optimal potency in the TTR-RBP4 HTRF assay (IC_50_ = 194 nM). AKR-XI-85 was selected as the lead compound for further derivatization.

**Table 2.**
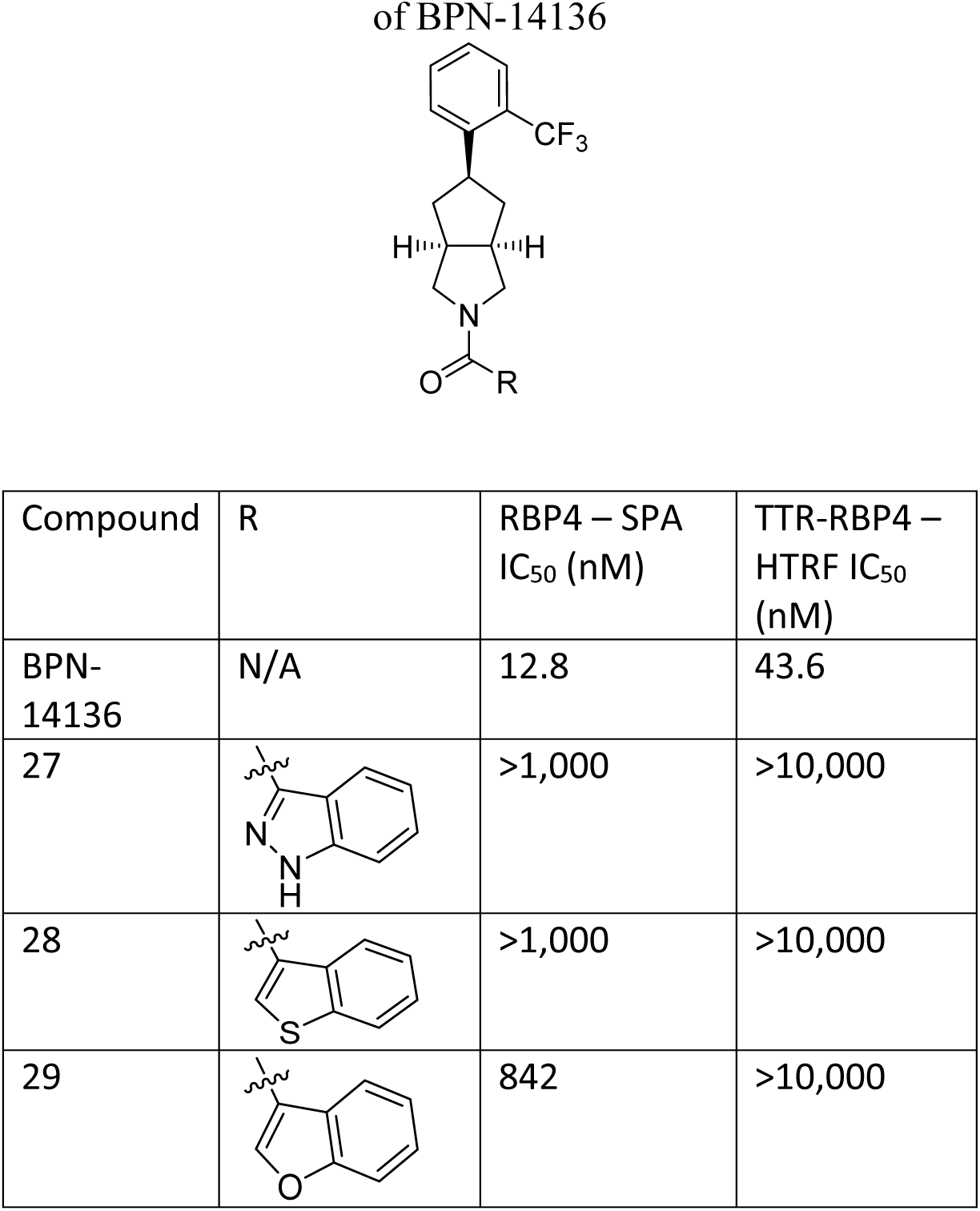
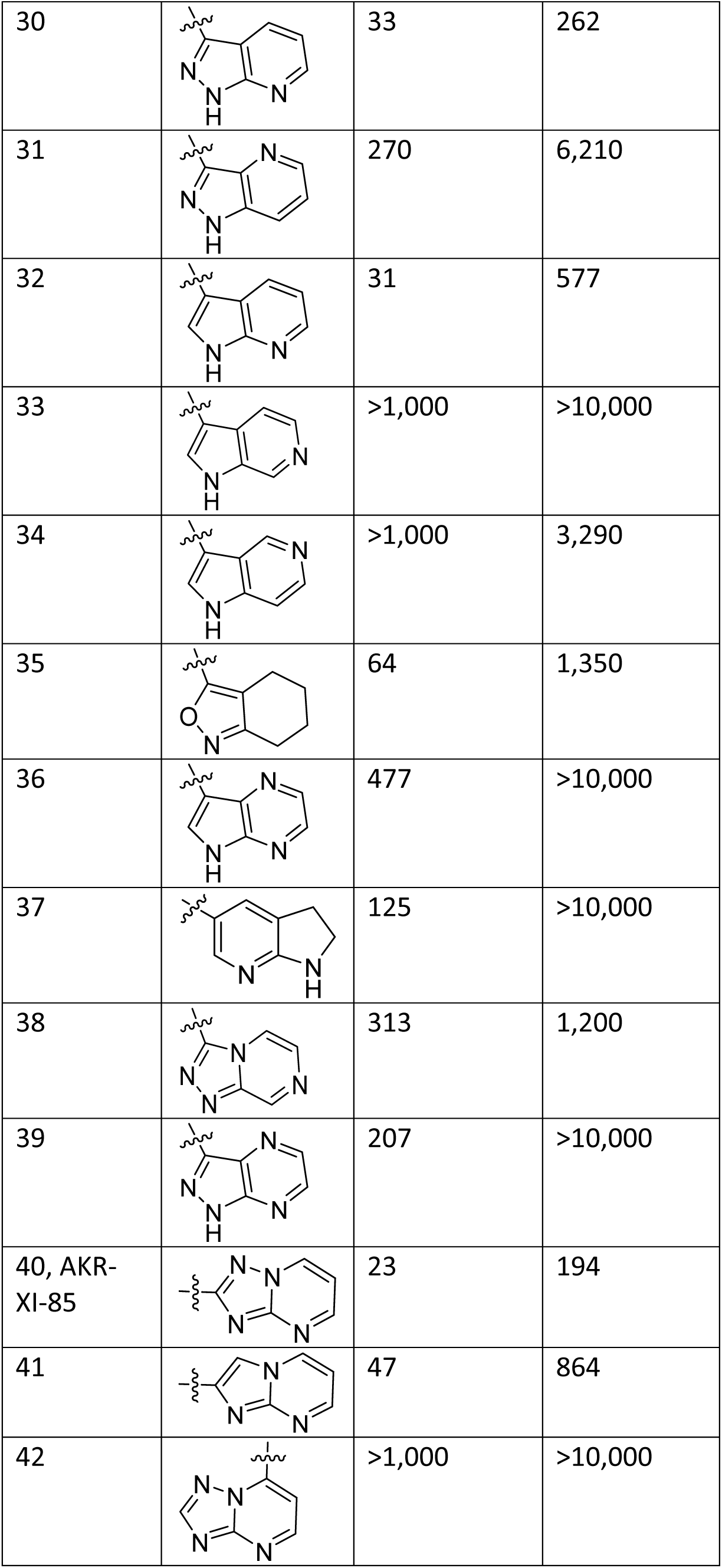
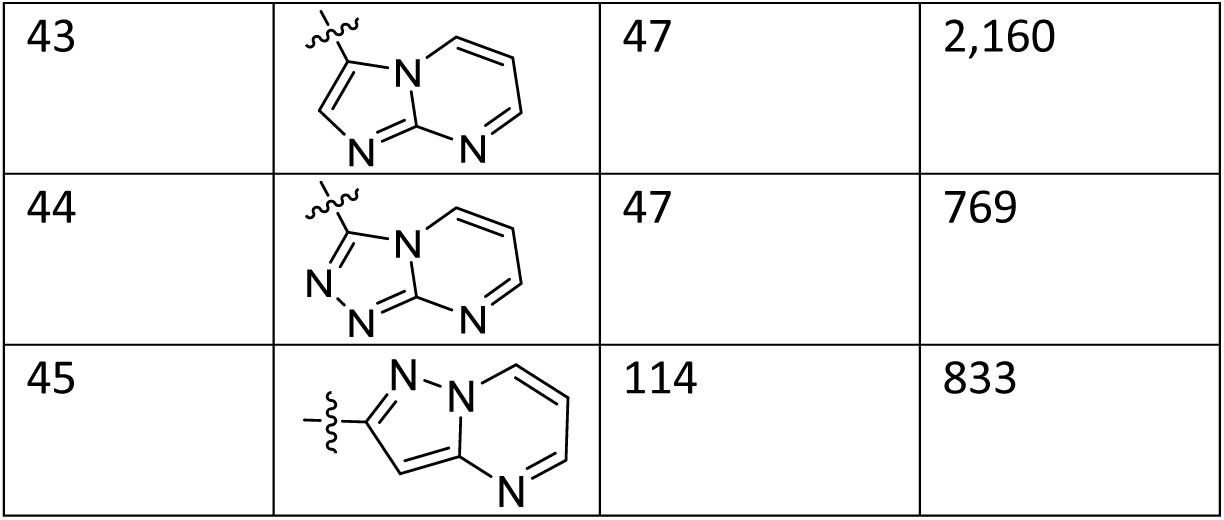
RBP4 binding affinity and functional RBP4-TTR antagonism for “bottom ring” analogs of BPN-14136.

| Compound | R | RBP4 – SPA<br>$IC_{50}$ (nM) | TTR-RBP4 –<br>HTRF $IC_{50}$<br>(nM) |
| --- | --- | --- | --- |
| BPN-14136 | N/A | 12.8 | 43.6 |
| 27 |  | >1,000 | >10,000 |
| 28 |  | >1,000 | >10,000 |
| 29 |  | 842 | >10,000 |
| 30 |  | 33 | 262 |
| 31 |  | 270 | 6,210 |
| 32 |  | 31 | 577 |
| 33 |  | >1,000 | >10,000 |
| 34 |  | >1,000 | 3,290 |
| 35 |  | 64 | 1,350 |
| 36 |  | 477 | >10,000 |
| 37 |  | 125 | >10,000 |
| 38 |  | 313 | 1,200 |
| 39 |  | 207 | >10,000 |
| 40, AKR-XI-85 |  | 23 | 194 |
| 41 |  | 47 | 864 |
| 42 |  | >1,000 | >10,000 |
| 43 |  | 47 | 2,160 |
| 44 |  | 47 | 769 |
| 45 |  | 114 | 833 |

Several derivatives of AKR-XI-85 were synthesized. First, we explored the effect of substituents on the triazolopyrimidine ring (Table 3). Introduction of small alkyl groups, such as methyl (**46, 49-50**) and fluorinated alkyl (**47, 51-52**) groups led to a reduction in RBP4 binding affinities and a dramatic loss in TTR-RBP4 efficacy, especially in analogs with a C-7 substituent. The C-6 vinyl analog (**62**) displayed moderate RBP4 binding (IC_50_ = 66 nM). While the C-6 propargyl alcohol (**79**) analog showed good binding (IC_50_ = 48 nM) but weak RBP4-TTR interaction potency, the prop-2-ynylphenyl (**80**) compound displayed no affinity for RBP4 and no RBP4-TTR interaction efficacy. Exceptional RBP4 binding was seen with a 2-hydroxyethyl group at the C-6 position (**61**, IC_50_ = 16 nM); however, the potency in the TTR-RBP4 HTRF assay was lower than that of AKR-XI-85 (IC_50_ = 246 nM). The methyl ether (**67**) and acetyl ester (**68**) and even the bulky tosylate (**65**) of **61** retained good RBP4 binding but **67** had poor potency in blocking the retinol-dependent TTR-RBP4 interaction. The C-6 carboxymethyl analog (**69**), along with its methyl ester (**70**) and amide (**71**) showed meager RBP4 binding and poor TTR-RBP4 interaction potency. The presence of an amino group either directly attached to the triazolopyrimidine ring at the C-7 position (**54**) or as part of a C-6 alkyl substituent (**63**, **66**, **72-78**) resulted in a significant reduction of RBP4 binding affinity and a decrease in TTR-RBP4 interaction potency ranging from 8-fold (**54**) to complete elimination (**73-74**). A select group of compounds (**48-50, 63, 64, 67, 68**) was tested in the PPARγ agonism assay where they showed no activity (IC_50_ > 100 µM).

**Table 3.**
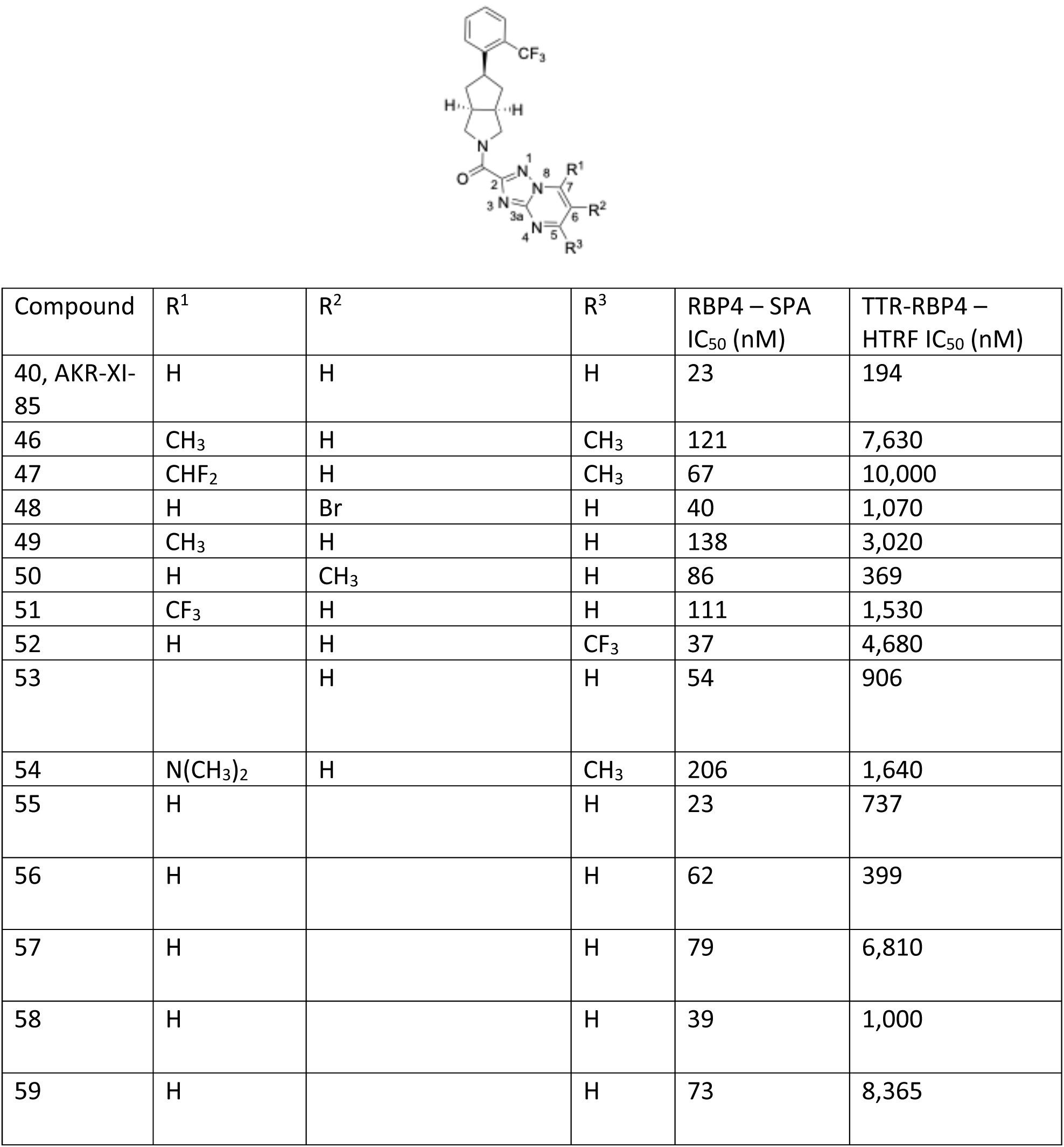
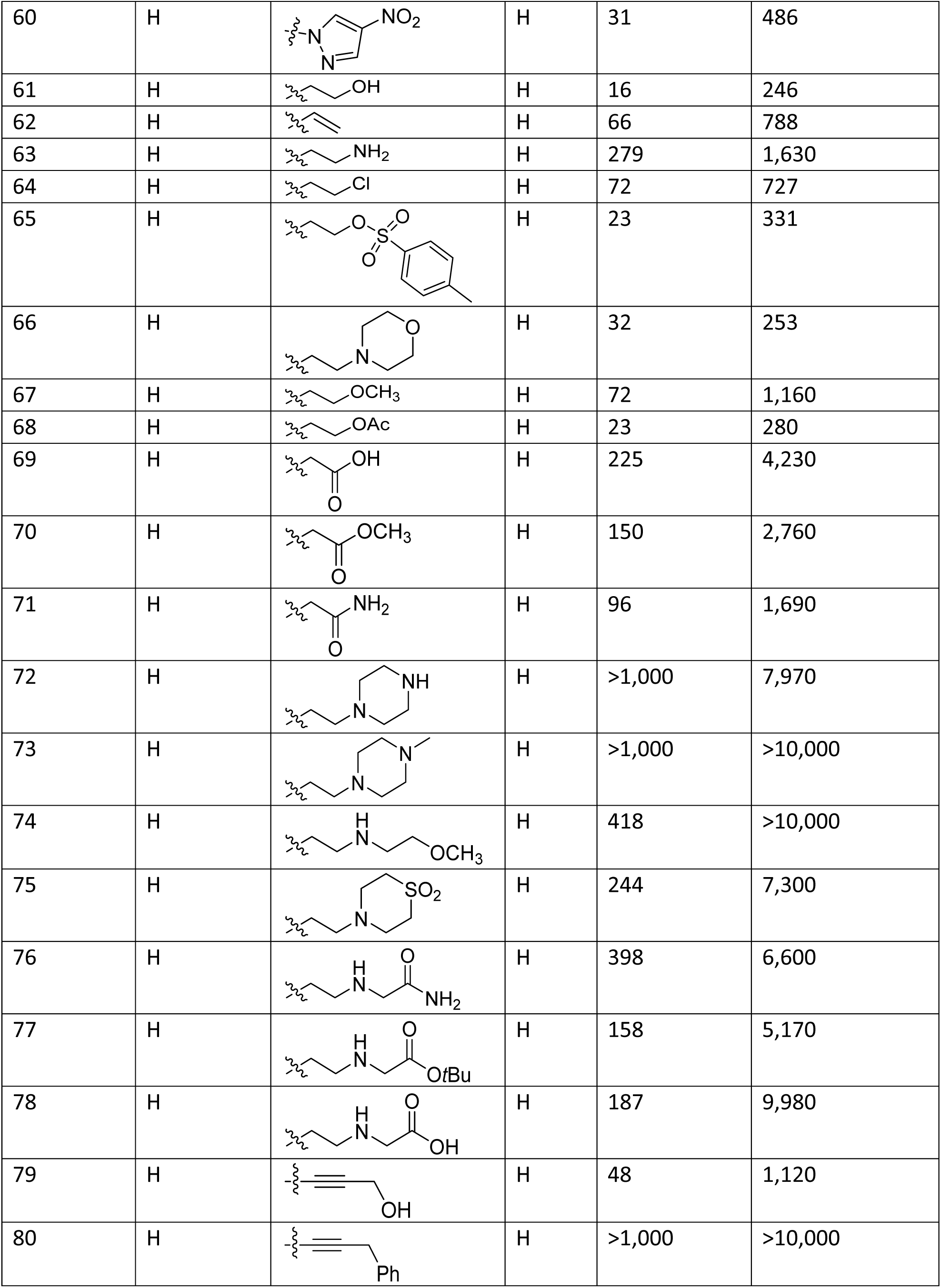
RBP4 binding affinity and functional RBP4-TTR antagonism for analogs of AKR-XI-85 (40) with triazolopyrimidine ring substituents.

| Compound | R<sup>1</sup> | R<sup>2</sup> | R<sup>3</sup> | RBP4 – SPA<br>IC<sub>50</sub> (nM) | TTR-RBP4 –<br>HTRF IC<sub>50</sub> (nM) |
| --- | --- | --- | --- | --- | --- |
| 40, AKR-XI-85 | H | H | H | 23 | 194 |
| 46 | CH<sub>3</sub> | H | CH<sub>3</sub> | 121 | 7,630 |
| 47 | CHF<sub>2</sub> | H | CH<sub>3</sub> | 67 | 10,000 |
| 48 | H | Br | H | 40 | 1,070 |
| 49 | CH<sub>3</sub> | H | H | 138 | 3,020 |
| 50 | H | CH<sub>3</sub> | H | 86 | 369 |
| 51 | CF<sub>3</sub> | H | H | 111 | 1,530 |
| 52 | H | H | CF<sub>3</sub> | 37 | 4,680 |
| 53 |  | H | H | 54 | 906 |
| 54 | N(CH<sub>3</sub>)<sub>2</sub> | H | CH<sub>3</sub> | 206 | 1,640 |
| 55 | H |  | H | 23 | 737 |
| 56 | H |  | H | 62 | 399 |
| 57 | H |  | H | 79 | 6,810 |
| 58 | H |  | H | 39 | 1,000 |
| 59 | H |  | H | 73 | 8,365 |
| 60 | H |  | H | 31 | 486 |
| 61 | H |  | H | 16 | 246 |
| 62 | H |  | H | 66 | 788 |
| 63 | H |  | H | 279 | 1,630 |
| 64 | H |  | H | 72 | 727 |
| 65 | H |  | H | 23 | 331 |
| 66 | H |  | H | 32 | 253 |
| 67 | H |  | H | 72 | 1,160 |
| 68 | H |  | H | 23 | 280 |
| 69 | H |  | H | 225 | 4,230 |
| 70 | H |  | H | 150 | 2,760 |
| 71 | H |  | H | 96 | 1,690 |
| 72 | H |  | H | >1,000 | 7,970 |
| 73 | H |  | H | >1,000 | >10,000 |
| 74 | H |  | H | 418 | >10,000 |
| 75 | H |  | H | 244 | 7,300 |
| 76 | H |  | H | 398 | 6,600 |
| 77 | H |  | H | 158 | 5,170 |
| 78 | H |  | H | 187 | 9,980 |
| 79 | H |  | H | 48 | 1,120 |
| 80 | H |  | H | >1,000 | >10,000 |

Next, we turned our attention to the head group. The 2-(trifluoromethyl)phenyl head group is a common structural feature in BPN-14136 and the archetypical RBP4 ligand that served as the starting point for its design, A1120. (19, 21, 22) We created a series of compounds to explore the effect of various polar and non-polar ring substituents at all available positions (Table 4). In addition, we also replaced the phenyl group with heterocyclic rings. The 2-trifluoromethoxy (**93**) derivative displayed very good RBP4 binding (IC_50_ = 41 nM) and moderate potency in the RBP4-TTR interaction assay (IC_50_ = 573 nM) while the 2-pentafluoroethyl analog (**95**) showed exceptional RBP4 binding (IC_50_ = 27 nM), albeit low potency (IC50 = 1,100 nM) in the RBP4-TTR interaction assay. Interestingly, the 2-isopropyl derivative (**94**) had a more favorable *in vitro* potency than the less bulky 2-methyl (**88**) and 2-ethyl (**91**) compounds. Polar substituents at the C-4 position (**96, 97**) were not tolerated. The introduction of heteroatoms into the head group ring completely eliminated all *in vitro* potency. Two high-affinity compounds, **82** and **93**, were tested in the PPARγ agonism assay where they showed no activity (IC_50_ > 100 µM).

**Table 4.**
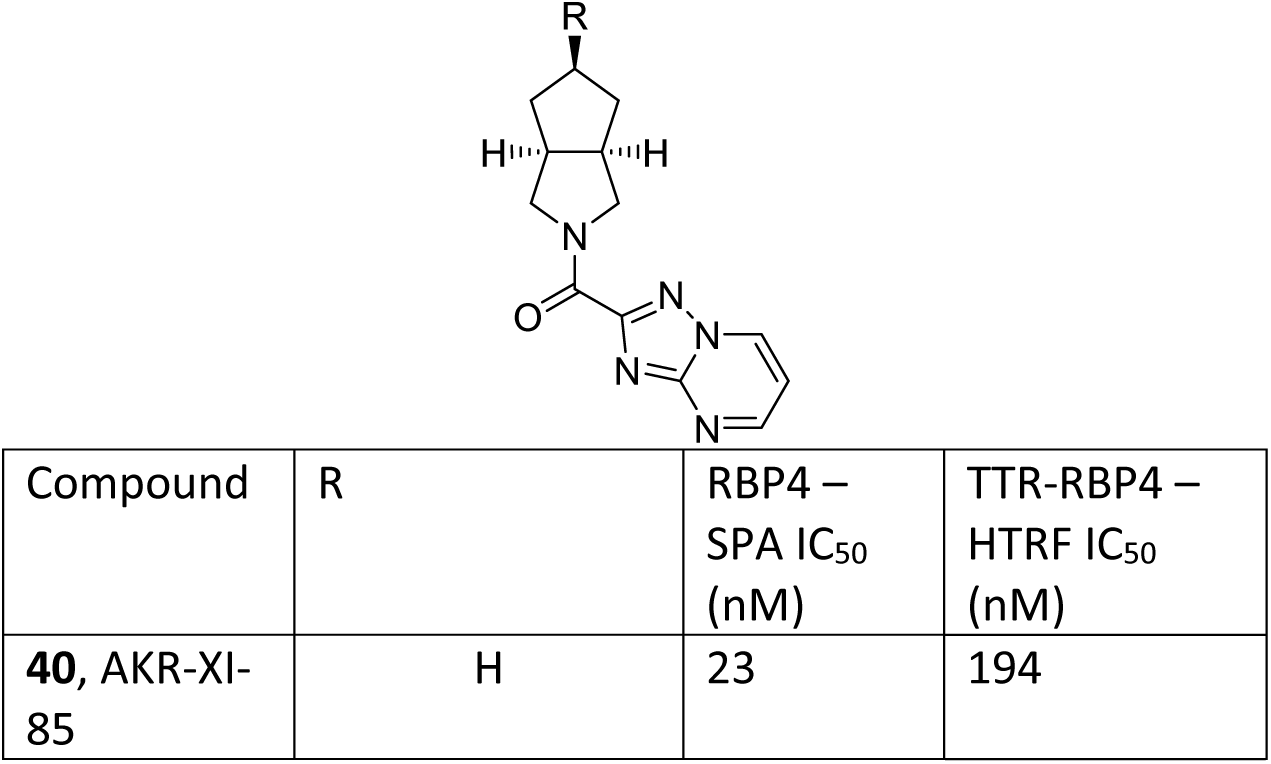
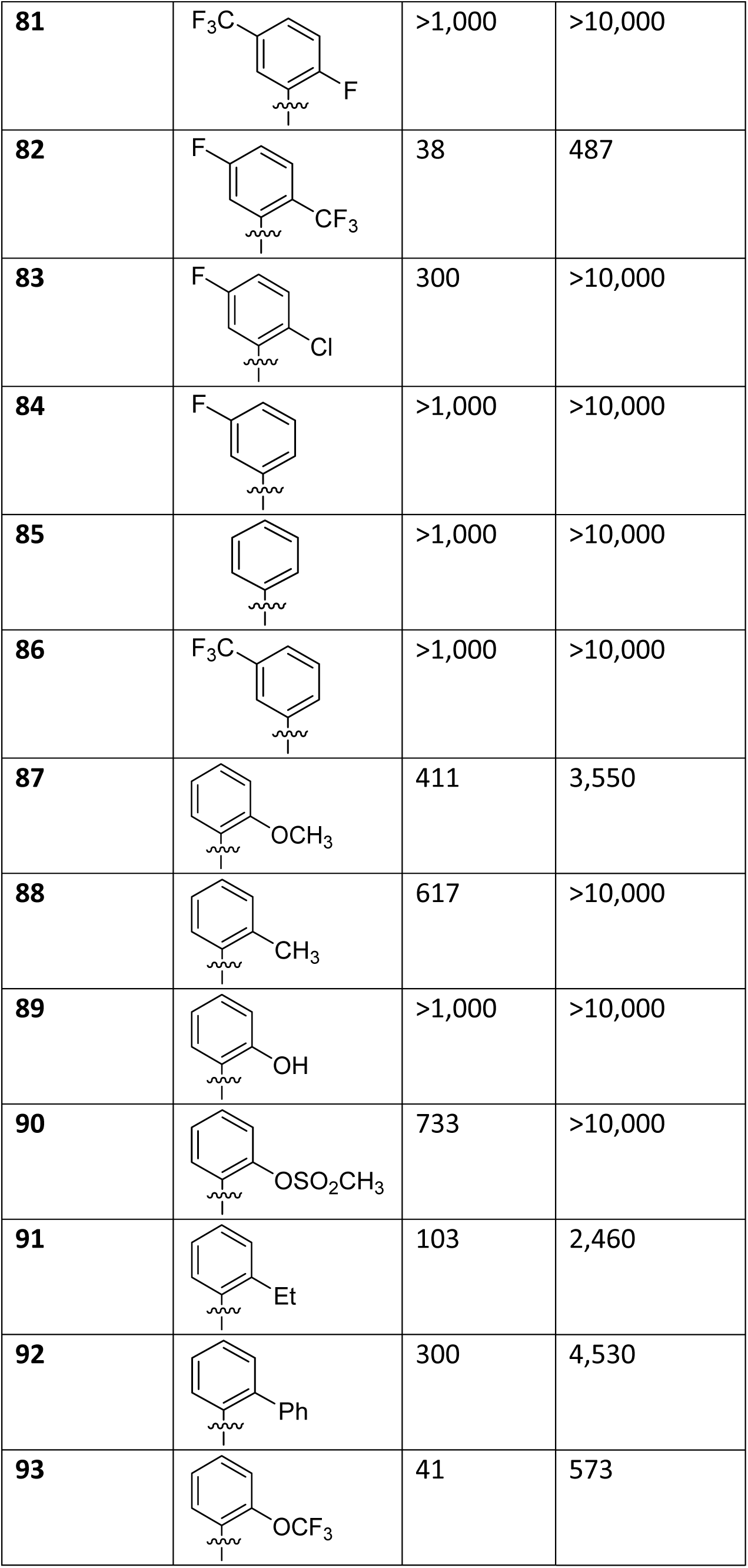
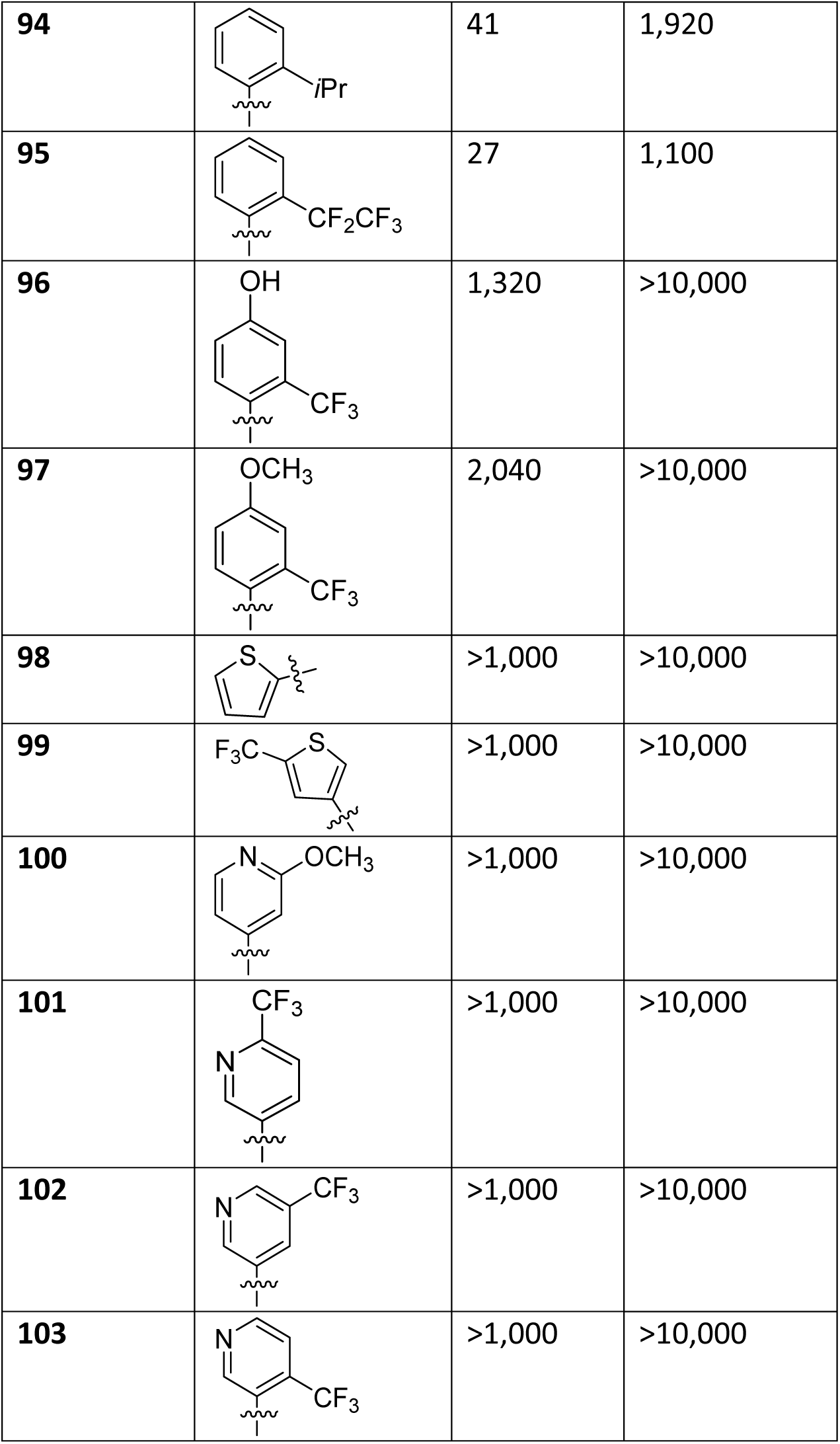
Analogs of AKR-XI-85 (40) with modified head groups.

| Compound | R | RBP4 –<br>SPA $IC_{50}$<br>(nM) | TTR-RBP4 –<br>HTRF $IC_{50}$<br>(nM) |
| --- | --- | --- | --- |
| **40**, AKR-XI-85 | H | 23 | 194 |
| 81 |  | >1,000 | >10,000 |
| 82 |  | 38 | 487 |
| 83 |  | 300 | >10,000 |
| 84 |  | >1,000 | >10,000 |
| 85 |  | >1,000 | >10,000 |
| 86 |  | >1,000 | >10,000 |
| 87 |  | 411 | 3,550 |
| 88 |  | 617 | >10,000 |
| 89 |  | >1,000 | >10,000 |
| 90 |  | 733 | >10,000 |
| 91 |  | 103 | 2,460 |
| 92 |  | 300 | 4,530 |
| 93 |  | 41 | 573 |
| 94 |  | 41 | 1,920 |
| 95 |  | 27 | 1,100 |
| 96 |  | 1,320 | >10,000 |
| 97 |  | 2,040 | >10,000 |
| 98 |  | >1,000 | >10,000 |
| 99 |  | >1,000 | >10,000 |
| 100 |  | >1,000 | >10,000 |
| 101 |  | >1,000 | >10,000 |
| 102 |  | >1,000 | >10,000 |
| 103 |  | >1,000 | >10,000 |

As a result of our SAR campaign, AKR-XI-85 (**40**) emerged as a lead due to its favorable *in vitro* potency in both the RBP4 binding and RBP4-TTR interaction assays, along with its absence of activity in the PPARγ agonism assay.

### 3.3 *In vitro* ADMET characteristics of AKR-XI-85

AKR-XI-85 was selected for further *in vitro* evaluation. The *in vitro* pharmacological profile is presented in Tables 5 and 6. As shown in Table 6, AKR-XI-85 exhibited excellent kinetic and very good equilibrium solubility in phosphate-buffered saline (PBS) (pH 7.4). The % plasma protein binding (PPB) data indicates a low fraction of unbound compound (Table 5). The observed high microsomal stability and CL_int_ values in dog and human liver microsomes suggest low predicted hepatic clearance in these two species. AKR-XI-85 demonstrated low metabolic stability in rat liver microsomes (Table 5), which could lead to limited exposure of the compound in rats. This may hinder the assessment of AKR-XI-85’s efficacy and safety in rats. However, the compound exhibited reasonable stability in mouse liver microsomes (Table 5), which supports its further evaluation in the mouse model of enhanced retinal lipofuscinogenesis. Additionally, adequate stability of AKR-XI-85 in mouse liver microsomes, is consistent with the selection of mice as a potential rodent safety species. We conducted a metabolite identification study following the incubation of AKR-XI-85 with human, rat, mouse, dog, and cynomolgus monkey liver microsomes. This study did not reveal any human-specific metabolites (data not shown), suggesting the feasibility of employing a standard set of rodent and non-rodent species in safety assessment studies for AKR-XI-85.

**Table 5.**
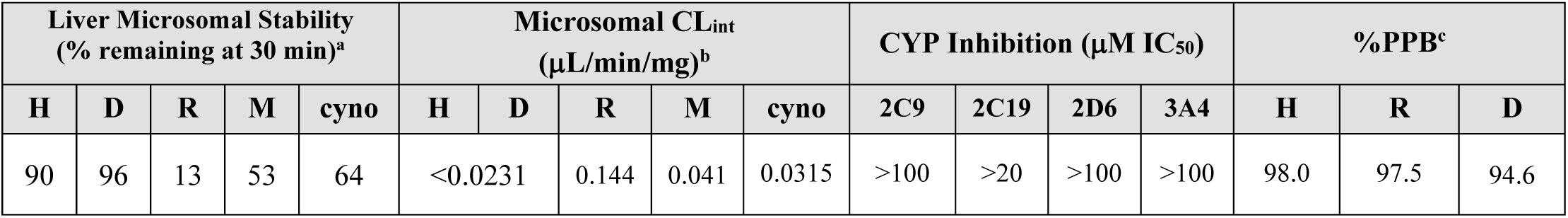
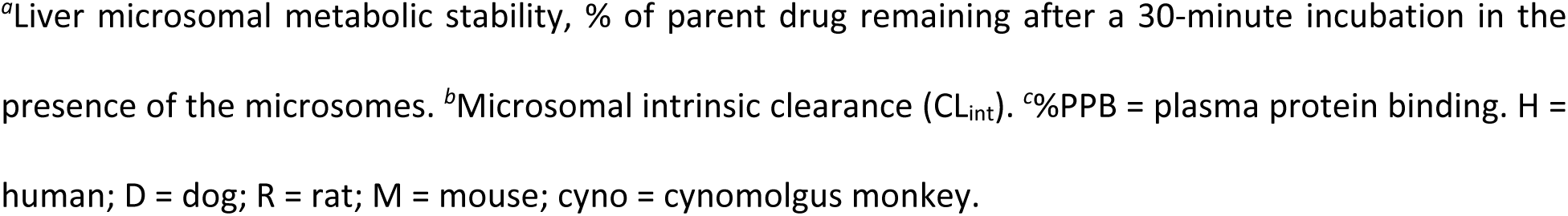
*In vitro* metabolic stability, CYP Inhibition and %PPB Profiles for AKR-XI-85.

| Liver Microsomal Stability<br>(% remaining at 30 min)<sup>a</sup> |  |  |  |  | Microsomal CL<sub>int</sub><br>(μL/min/mg)<sup>b</sup> |  |  |  |  | CYP Inhibition (μM IC<sub>50</sub>) |  |  |  | %PPB<sup>c</sup> |  |  |
| --- | --- | --- | --- | --- | --- | --- | --- | --- | --- | --- | --- | --- | --- | --- | --- | --- |
| H | D | R | M | cyno | H | D | R | M | cyno | 2C9 | 2C19 | 2D6 | 3A4 | H | R | D |
| 90 | 96 | 13 | 53 | 64 | <0.0231 |  | 0.144 | 0.041 | 0.0315 | >100 | >20 | >100 | >100 | 98.0 | 97.5 | 94.6 |

<sup>a</sup>Liver microsomal metabolic stability, % of parent drug remaining after a 30-minute incubation in the presence of the microsomes. <sup>b</sup>Microsomal intrinsic clearance (CL<sub>int</sub>). <sup>c</sup>%PPB = plasma protein binding. H = human; D = dog; R = rat; M = mouse; cyno = cynomolgus monkey.

**Table 6.**
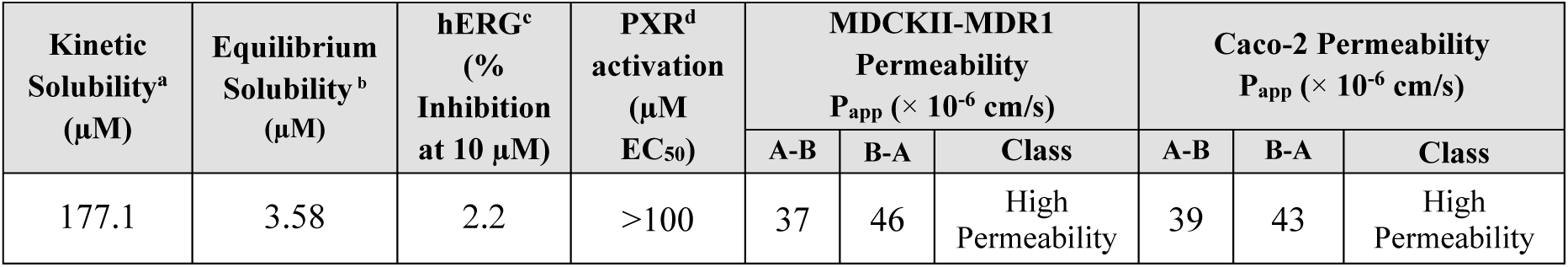
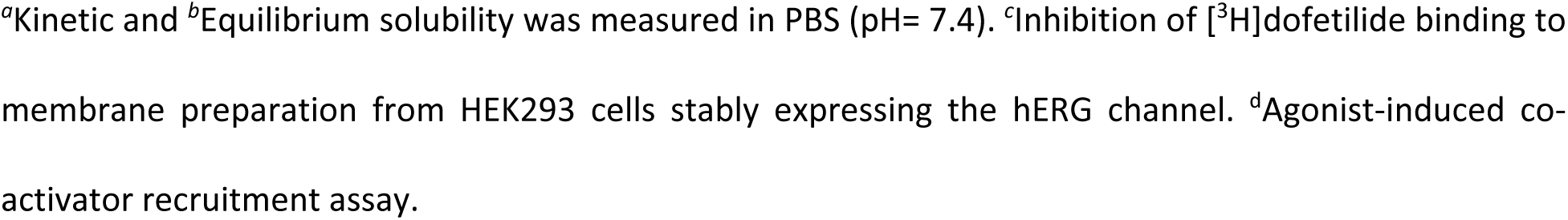
*In vitro* solubility, hERG inhibition, PXR activation and permeability profiles for AKR-XI-85.

| Kinetic Solubility<sup>a</sup><br>(μM) | Equilibrium Solubility<sup>b</sup><br>(μM) | hERG<sup>c</sup><br>(%<br>Inhibition<br>at 10 μM) | PXR<sup>d</sup><br>activation<br>(μM<br>EC<sub>50</sub>) | MDCKII-MDR1<br>Permeability<br>P<sub>app</sub> (× 10<sup>-6</sup> cm/s) |  |  | Caco-2 Permeability<br>P<sub>app</sub> (× 10<sup>-6</sup> cm/s) |  |  |
| --- | --- | --- | --- | --- | --- | --- | --- | --- | --- |
|  |  |  |  | A-B | B-A | Class | A-B | B-A | Class |
| 177.1 | 3.58 | 2.2 | >100 | 37 | 46 | High<br>Permeability | 39 | 43 | High<br>Permeability |

<sup>a</sup>Kinetic and <sup>b</sup>Equilibrium solubility was measured in PBS (pH= 7.4). <sup>c</sup>Inhibition of [<sup>3</sup>H]dofetilide binding to membrane preparation from HEK293 cells stably expressing the hERG channel. <sup>d</sup>Agonist-induced co-activator recruitment assay.

AKR-XI-85 lacked limiting inhibitory activity in a standard CYP 450 panel (Table 5). Additional selectivity profiling revealed no significant off-target activity at the hERG channel (Table 6) or within a standard Cerep-Panlabs SafetyScreen 44 panel of forty-four GPCRs, enzymes, ion channels, and transporters (data not shown). Lastly, AKR-XI-85 did not induce human PXR activation (Table 6). The results of permeability testing in MDCKII-MDR1 and Caco-2 cells (Table 6) indicate that AKR-XI-85 is a highly permeable compound that is expected to have good *in vivo* absorption. The compound does not appear to be the substrate for the Pgp efflux transporter. To evaluate the potential mutagenic and genotoxic properties of AKR-XI-85 we performed its characterization in Ames studies using the following strains of *Salmonella typhimurium*: TA98, TA100, TA1535, and TA1537. The experiments were conducted in the presence and absence of the microsomal S9 fraction. AKR-XI-85 showed no signs of genotoxicity and mutagenicity in the Ames studies (data not shown).

### 3.4 Pharmacokinetic (PK) characteristics of AKR-XI-85

Based on the outcomes of *in vitro* ADMET studies, mouse and dog were selected as the two species for performing single-dose PK studies. The PK properties of AKR-XI-85 following a single oral (5 mg/kg) and intravenous (2 mg/kg) dose are shown in Table 7. In mice, the compound showed adequate bioavailability (59.6 %) and rapid clearance following both routes of administration. Due to rapid systemic clearance, a rather limited exposure was achieved in mice. However, as demonstrated below, the PK properties of AKR-XI-85 in mice were sufficient to achieve the desired pharmacodynamic effect in this species (robust RBP4 reduction) and attain the intended bisretinoid-lowering efficacy in the mouse *Abca4^-/-^* model. In dogs, however, AKR-XI-85 displayed low clearance and a high exposure after both oral and intravenous administration (AUC_INF_ = 15,698 h*ng/ml and 27,593 h*ng/ml, respectively), which is in line with the favorable metabolic stability observed in the dog liver microsomal experiment. The favorable PK profile of AKR-XI-85 in dogs, along with the comparatively lower exposure observed in mice, emphasizes the significance of exploring the pharmacodynamic response in mice (serum RBP4 reduction), given that RBP4 lowering directly determines the efficacy of RBP4 antagonists in reducing lipofuscin bisretinoids in the retina.

**Table 7.**
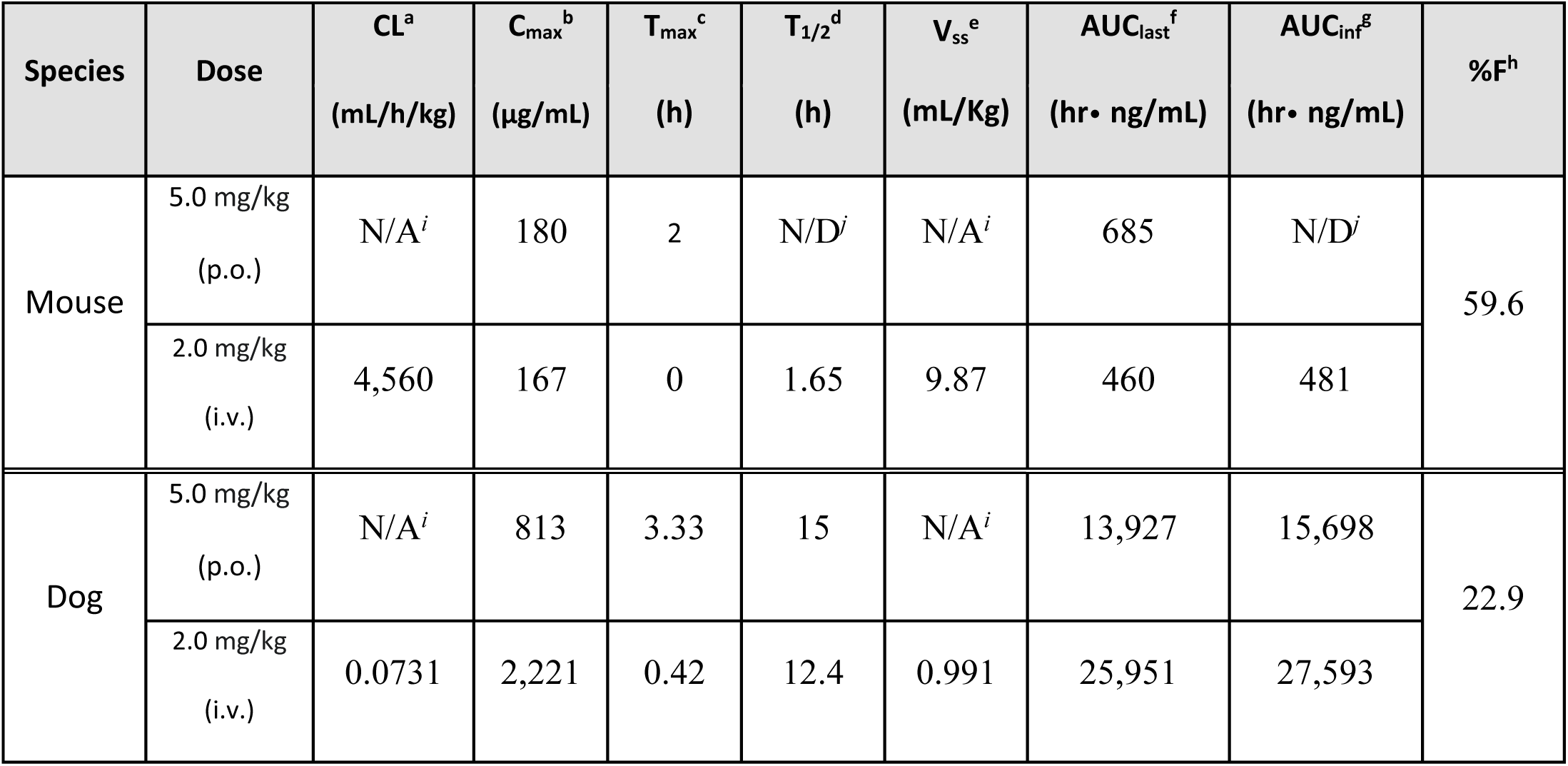
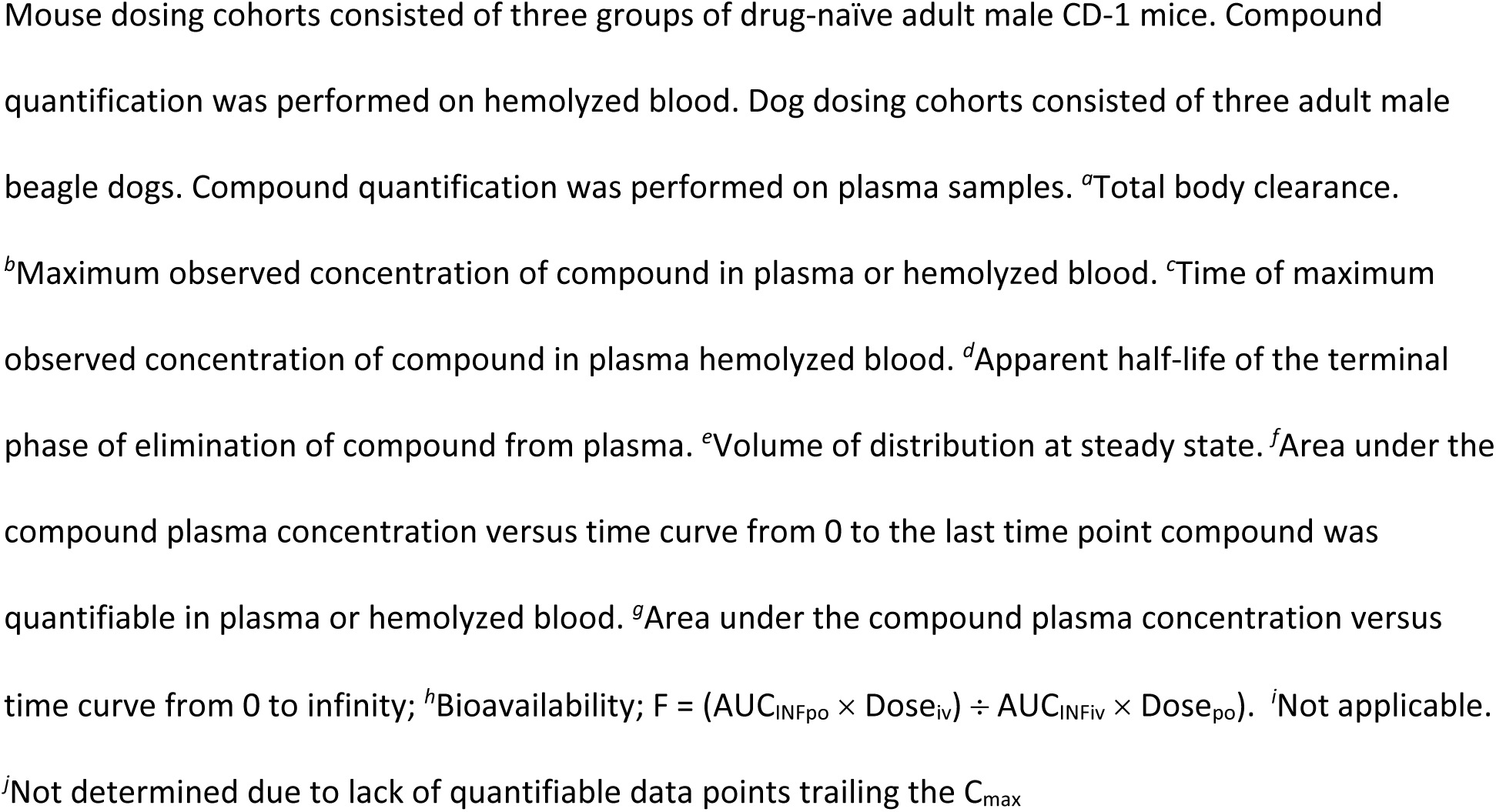
*In vivo* PK data for AKR-XI-85 following intravenous (i.v.) and oral (p.o.) administration in mouse and beagle dog.

| Species | Dose | CL<sup>a</sup><br>(mL/h/kg) | C<sub>max</sub><sup>b</sup><br>(µg/mL) | T<sub>max</sub><sup>c</sup><br>(h) | T<sub>1/2</sub><sup>d</sup><br>(h) | V<sub>ss</sub><sup>e</sup><br>(mL/Kg) | AUC<sub>last</sub><sup>f</sup><br>(hr• ng/mL) | AUC<sub>inf</sub><sup>g</sup><br>(hr• ng/mL) | %F<sup>h</sup> |
| --- | --- | --- | --- | --- | --- | --- | --- | --- | --- |
| Mouse | 5.0 mg/kg<br>(p.o.) | N/A<sup>i</sup> | 180 | 2 | N/D<sup>j</sup> | N/A<sup>i</sup> | 685 | N/D<sup>j</sup> | 59.6 |
|  | 2.0 mg/kg<br>(i.v.) | 4,560 | 167 | 0 | 1.65 | 9.87 | 460 | 481 |  |
| Dog | 5.0 mg/kg<br>(p.o.) | N/A<sup>i</sup> | 813 | 3.33 | 15 | N/A<sup>i</sup> | 13,927 | 15,698 | 22.9 |
|  | 2.0 mg/kg<br>(i.v.) | 0.0731 | 2,221 | 0.42 | 12.4 | 0.991 | 25,951 | 27,593 |  |

Mouse dosing cohorts consisted of three groups of drug-naïve adult male CD-1 mice. Compound quantification was performed on hemolyzed blood. Dog dosing cohorts consisted of three adult male beagle dogs. Compound quantification was performed on plasma samples. <sup>a</sup>Total body clearance. <sup>b</sup>Maximum observed concentration of compound in plasma or hemolyzed blood. <sup>c</sup>Time of maximum observed concentration of compound in plasma hemolyzed blood. <sup>d</sup>Apparent half-life of the terminal phase of elimination of compound from plasma. <sup>e</sup>Volume of distribution at steady state. <sup>f</sup>Area under the compound plasma concentration versus time curve from 0 to the last time point compound was quantifiable in plasma or hemolyzed blood. <sup>g</sup>Area under the compound plasma concentration versus time curve from 0 to infinity; <sup>h</sup>Bioavailability; $F = (AUC_{INFpo} \times Dose_{iv}) \div AUC_{INFiv} \times Dose_{po}$ . <sup>i</sup>Not applicable. <sup>j</sup>Not determined due to lack of quantifiable data points trailing the C<sub>max</sub>

### 3.5 Pharmacodynamics of AKR-XI-85 in mice

The interaction between RBP4 and TTR depends on retinol binding to RBP4. When an RBP4 antagonist displaces retinol from serum RBP4, it triggers the dissociation of the circulating RBP4-TTR complex. This, in turn, leads to the renal clearance of RBP4 from the bloodstream due to the small size of apo-RBP4 released from the complex with TTR. Assessing serum RBP4 concentrations provides a convenient pharmacodynamics (PD) marker for evaluating the target engagement and *in vivo* potency of RBP4 antagonists. We studied the effect of single escalating doses of AKR-XI-85 on the concentration of serum RBP4 in mice. Serum samples collected after oral administration of AKR-XI-85 were used to analyze RBP4 concentrations in the ELISA assay. After a single 15 mg kg^−1^ oral dose of AKR-XI-85, administered through oral gavage, a maximum of an 83% decrease in serum RBP4 was observed at 4 hours after dosing. A single 25 mg kg^−1^ oral dose induced a maximum of an 86% decrease in serum RBP4 at 6 hours post-dosing, while the 35 mg kg^−1^ oral dose resulted in a maximum of an 89% reduction at 8 hours after compound administration (Fig. 4). The effect of dose escalation on duration of the RBP4 lowering effect was also evident. After a single 15 mg kg^−1^ dose, there was a complete rebound of serum RBP4 concentration at the 24-h time point (Fig. 4). However, for doses of 25 mg kg^−1^ and 35 mg kg^−1^, a 5% and a 25% reduction in RBP4, respectively, was still observed at the 24-h time point. Overall, our data strongly supports a consistently positive biological PD response to AKR-XI-85 in mice.

**Fig 4.**
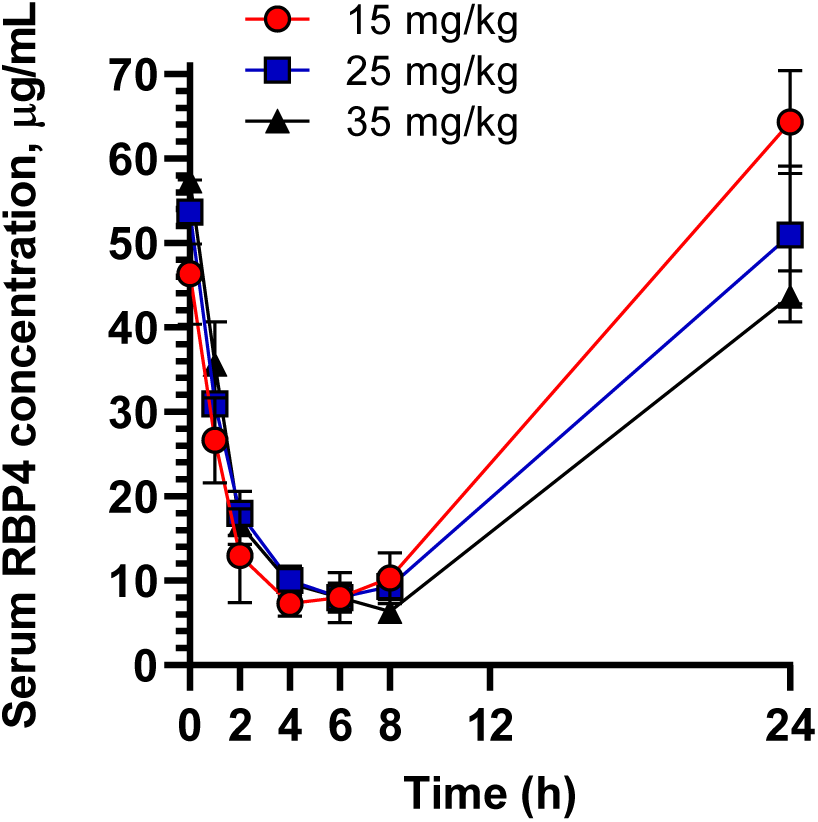
Pharmacodynamic properties of AKR-XI-85 in mice. Serum RBP4 levels in Balb/c mice following a single oral administration of AKR-XI-85 at a dose of 15 mg kg^−1^(red), 25 mg kg^−1^ (blue), and 35 mg kg^−1^ (black). Compound was formulated in 2% Tween 80 and 0.9% saline. Data are represented as the mean ± S.D. Three mice per time point of blood collection were used.

### 3.6 *In vivo* efficacy of AKR-XI-85 in the *Abca4^-/-^*mouse model of Stargardt disease

The *Abca4 ^-/-^* mouse model is widely regarded as the gold standard for assessing the preclinical efficacy of bisretinoid-lowering therapies. (39–43) To assess the ability of AKR-XI-85 to reduce bisretinoid levels in the retina, AKR-XI-85 was orally administered to *Abca4^-/-^* mice at a daily dose of 35 mg kg^−1^ for a duration of 60 days. The compound was incorporated into the chow to facilitate consistent oral dosing throughout the duration of the treatment. Blood samples were collected from treatment groups at baseline and again at the end of the dosing period to measure the level of serum RBP4. Chronic treatment of *Abca4^-/-^* mice with AKR-XI-85 resulted in a significant reduction in serum RBP4 levels, while no RBP4 reduction was observed in the vehicle-treated *Abca4^-/-^*mice. (Fig. 5A). At the end of the treatment period, compound-treated *Abca4^-/-^* mice exhibited a 72% reduction in RBP4 levels compared to untreated animals. After the end of the treatment period, levels of lipofuscin fluorophores (A2E) were quantified using HPLC in eyecup extracts. Long-term oral dosing of AKR-XI-85 induced a robust, 70% reduction in the A2E concentration in the retinas of *Abca4^-/-^* mice compared with vehicle-treated animals of identical genetic background (Fig. 5B). Prolonged exposure to AKR-XI-85 did not result in any observable toxicity and had no effect on the body weight or food intake of the animals, suggesting a lack of significant toxicity.

**Fig. 5.**
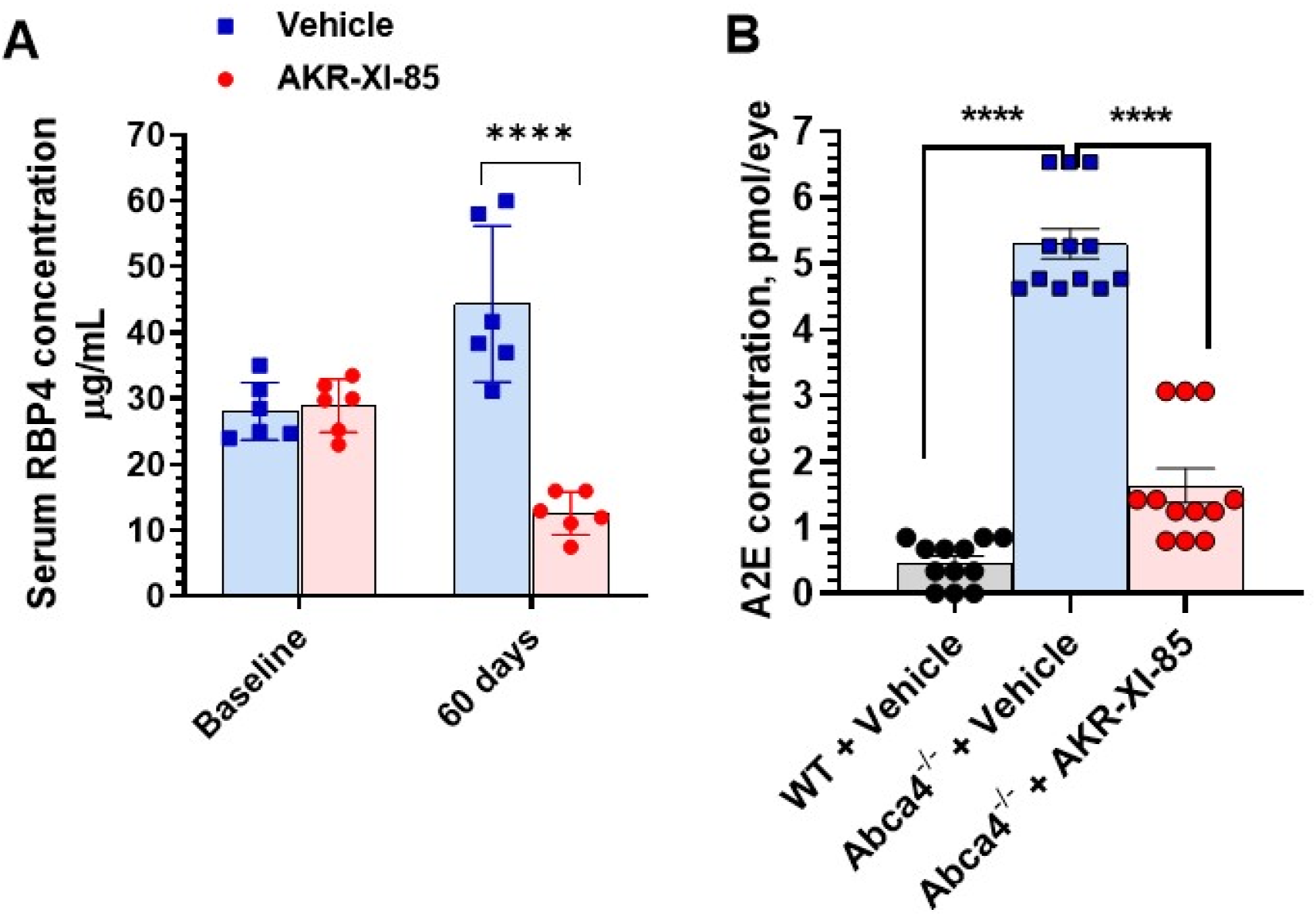
*In vivo* efficacy of AKR-XI-85 in the *Abca4^-/-^* model of Stargardt disease. ***<u>A</u>***, Serum RBP4 levels were measured in vehicle-treated *Abca4^-/-^*mice (blue squares; n=6) and AKR-XI-85-treated *Abca4^-/-^* mice (red circles, n=6) at the indicated time points. AKR-XI-85 was formulated into the chow and was dosed at 35 mg kg^−1^. Compared with the vehicle-treated group, a statistically significant RBP4 reduction was seen upon the termination of the experiment in the AKR-XI-85 treatment group (multiple t-test; discovery determined using the two-stage linear step-up procedure of Benjamini, Krieger, and Yekutieli. Each row was analyzed individually, without assuming a consistent S.D., p < 0.001). Error bars show S.D. Each data point on the graph represents a serum RBP4 concentration from an individual animal. ***<u>B</u>***, Effect of AKR-XI-85 treatment on A2E levels in the eyes of *Abca4^−/−^* mice. Bisretinoids were extracted from the eyecups of vehicle-treated *Abca4^+/+^* (wild-type, WT) mice (black circles), vehicle-treated *Abca4^−/−^* mice (blue squares), and AKR-XI-85-treated *Abca4^−/−^* (red circles) after 60 days of dosing. The A2E concentration in vehicle-treated WT mice was significantly lower than in vehicle-treated *Abca4^−/−^* KO animals (One-way ANOVA with Holm-Sidak post-hoc test, P < 0.0001). A significant, 70% reduction in the A2E levels was measured in AKR-XI-85-treated *Abca4^−/−^* mice in comparison to vehicle-treated knockout controls (One-way ANOVA with Holm-Sidak post-hoc test, P < 0.0001).

## 4. Discussion

Light perception in the retina (phototransduction) is initiated by the light-induced isomerization of the visual pigment, 11-*cis*-retinaldehyde bound to opsin, to the all-*trans* isomer. After the release from opsin, all-*trans*-retinal reacts with phosphatidylethanolamine through a reversible Schiff base, forming *N*-retinylidene-phosphatidylethanolamine (*N*-retinylidene-PE). The ABCA4 transporter expressed in photoreceptor cells flips *N*-retinylidene-PE across the disk membrane to the cytoplasmic side where the highly reactive all-*trans*-retinaldehyde can be released and reduced to the inactive all-*trans*-retinol. The prompt translocation of *N*-retinylidene-PE across the disk membrane by the ABCA4 transporter prevents the reaction between *N*-retinylidene-PE and another molecule of all-*trans* retinaldehyde. This reaction between *N*-retinylidene-PE and the second molecule of all-*trans* retinaldehyde, known as retinaldehyde dimerization, represents the first step in the formation and deleterious buildup of toxic lipofuscin bisretinoids, such as A2E. In Stargardt disease, the impaired function of the ABCA4 transporter slows down the conversion of highly reactive all-*trans*-retinaldehyde to inactive all-*trans*-retinol. This increases the chance of retinaldehyde dimerization and facilitates bisretinoid formation. Furthermore, ABCA4 dysfunction may lead to increased exposure of retinal structures to free retinaldehyde which is proposed to have an independent role in the pathogenesis of macular degeneration due to inherent retinaldehyde toxicity (44, 45). STGD1 is the most common form of juvenile-onset retinal degeneration caused by recessive mutations in the ABCA4 gene. The primary biochemical defect in STGD1 is the overproduction of cytotoxic lipofuscin bisretinoids in the retinal pigment epithelium (46, 47) caused by the malfunctioning ABCA4 transporter. With no FDA-approved therapies available, the treatment of STGD1 remains a critical unmet medical need. Given that cytotoxic lipofuscin bisretinoids are formed in the retina in a non-enzymatic fashion from visual cycle retinaldehydes (11, 48) several pharmacological strategies emerged to interfere with the formation of lipofuscin bisretinoids to slow or arrest the progression of Stargardt disease. (12, 49) Various classes of small-molecule drugs have been suggested to function as inhibitors of bisretinoid synthesis and mitigators of retinaldehyde toxicity, including RPE65 inhibitors, selective RBP4 antagonists, aldehyde traps, and deuterated analogs of vitamin A. (12, 33, 49, 50) Two classes of small-molecule inhibitors of bisretinoid formation which emerged recently are bispecific RBP4/TTR ligands and selective TTR ligands.(33, 50) AKR-XI-85 identified in this study as a potential therapy for Stargardt disease belongs to the class of selective RBP4 antagonists which inhibit bisretinoid synthesis by reducing the overall retinoid load on the retina.(51) The majority of retinol is stored in the liver from where it is carried to the target organs by the most abundant retinol transporter, RBP4. Retinol-bound RBP4 is stabilized by binding to TTR, while RBP4 devoid of retinol is unable to bind TTR and, in the absence of the TTR homotetramer, RBP4 is rapidly cleared from circulation. Therefore, RBP4 antagonists that displace retinol from RBP4 can induce dissociation of the circulating RBP4-TTR-retinol complex and reduce the levels of circulating RBP4 in the blood. Consequently, this diminishes the quantity of retinol transported to the retina and inhibits bisretinoid synthesis. Fenretinide, a synthetic retinoid drug originally developed for the treatment of cancer, can bind to RBP4 outcompeting retinol from its binding pocket. In a murine model of ABCA4-related retinopathies (*Abca4^-/-^*), fenretinide effectively reduced the levels of circulating RBP4 and diminished the formation of A2E in the eye.(42) In a clinical trial, orally given fenretinide slowed the progression of geographic atrophy in late-stage age-related macular degeneration (AMD).(52) Correlation between reduced lesion growth rates and serum RBP4 levels could be established at RBP4 levels of less than 2mg/dL. However, this level of RBP4 reduction was achieved in only 51% of the patients receiving the high-end dose of the therapeutic (300 mg/day), whereas no statistically significant effect could be determined in the lower dose group (100 mg/day). While the results are encouraging, there are concerns regarding the safety of long-term exposure to fenretinide as it can lead to apoptosis in the RPE(53) and, similar to other synthetic retinoids, may be teratogenic(54) and carcinogenic(55). While the clinical and preclinical efficacy of fenretinide firmly established the class of RBP4 antagonists as a potential therapy for macular degeneration, the development of non-retinoid RBP4 ligands was strongly desired to mitigate the safety concerns associated with retinoid RBP4 antagonists.

The first non-retinoid RBP4 ligand, A1120, was initially designed as a potential treatment for Type 2 diabetes. (19) However, despite its success in reducing serum RBP4 levels in mice, it did not result in any improvement in insulin resistance. (19) Furthermore, A1120 displayed low stability in human liver microsomes (19), rendering its development as a therapy for human diseases unfeasible. However, testing in mice was possible, and we showed that A1120 was effective in reducing the formation of toxic bisretinoids in the *Abca4^-/-^* mouse model of Stargardt disease. (20) Our subsequent efforts led to the discovery of several groups of promising non-retinoid compounds (21, 22, 24), including BPN-14136, an RBP4 antagonist with a bicyclic [3.3.0]-octahydrocyclopenta[*c*]pyrrolo core.(22, 23, 25) BPN-14136 exhibited several desirable properties, such as good *in vitro* potency, favorable PK/PD characteristics, remarkable lowering of circulating RBP4 levels *in vivo*, and a robust, approximately 50% reduction in the accumulation of cytotoxic bisretinoids in the eyes of *Abca4^-/-^* mice.(22, 23, 25) However, as we report here, BPN-14136 has an off-target activity as a PPARγ agonist. PPARγ is a nuclear receptor that is involved in maintaining the energy homeostasis and in regulating lipid biosynthesis. Agonistic activation of PPARγ is associated with increased risk of death, myocardial infarction, stroke, congestive heart failure, hepatotoxicity, peripheral edema, weight gain and carcinogenicity. (29–32) All known naturally occurring ligands of PPARγ are lipophilic carboxylic acids (56) and several synthetic ligands of PPARγ also share common structural features including a hydrophobic tail and a polar head group carrying a carboxylic acid group.(57–60) The heteroaryl carboxylic acid functional group featured in BPN-14136 seems to be involved in modulating the off-target activity of BPN-14136 as a PPARγ agonist (Fig. 3). Furthermore, the carboxylic acid featured in the BPN-14136 series may represent one additional liability. While the carboxylic acid group is a common appendage found on drug molecules that confers desirable pharmacological properties, such as reduced CNS permeability, this functional group has been linked to idiosyncratic drug toxicity, possibly due to the formation of reactive acyl glucuronide metabolites.(26–28) In this study, our aim was to design analogs of BPN-14136 devoid of the heteroaryl carboxylic functional group. Initially, two series of analogs were synthesized. First, we explored the effect of removing the carboxylic acid group and introducing different substituents on the bottom ring of BPN-14136 (Table 1). Overall, the substitution of the pyrimidine ring alone did not result in improved *in vitro* potency, but the absence of the carboxylic acid group resulted in the lack of PPARγ binding (IC_50_ > 100 µM). Next, the pyrimidine ring was replaced with a series of heterocycles and, to further reduce the risk of PPARγ engagement, a carbonyl group was also introduced between the bottom heterocycle and the [3.3.0]-octahydrocyclopenta[*c*]pyrrolo linker (Table 2). In this series, several analogs with high RBP4 affinity were identified (**30, 32, 35, 40, 41, 43, 44**). Compound **40**, AKR-XI-85, was chosen for further derivatization and analysis owing to its exceptional *in vitro* potency in antagonizing the TTR-RBP4 interaction (IC_50_ = 194 nM). Key binding parameters were revealed, using *in silico* docking studies (Fig.6). Similar to BPN-14136 (22), AKR-XI-85 occupies the binding pocket of RBP4 with the hydrophobic trifluoromethylphenyl group situated in the RBP4 interior and the triazolopyrimidine bottom ring is oriented toward and exposed to the solvent. The key interactions between the molecule and RBP4 include hydrogen bonding to Arg^121^. Groups with a size similar to that of the trifluoromethylphenyl group, such as 5-membered heterocyclic rings, a 2-methylphenyl, or a 2-trifluoromethoxyphenyl group, were tolerated in docking studies, whereas bulkier replacements resulted in significantly less favorable docking scores owing to steric clashes with the binding pocket. Substituents as large as a phenyl group can be accommodated at the C-6 position. Negatively charged appendages can also be accommodated, while the few positively-charged substituents modeled appear to be excluded. Encouraged by the *in silico* docking results, we synthesized two additional series of analogs using AKR-XI-85 as the template. The first series of derivatives explored the effect of substitutions on the triazolopyrimidine ring system (Table 3). C-7 substitutions resulted in reduced RBP4 affinity and a marked loss of activity in the retinol-dependent RBP4-TTR interaction assay. As expected from the docking studies, C-6 substitutions were better tolerated and good binding was seen with several analogs even with bulky substituents, such as an ethylene tosylate and a 2-*N*-morpholinoethylene group (**65, 66**). Overall, substitution of the triazolopyrimidine ring significantly reduced the ability of the compounds to antagonize the retinol-dependent TTR-RBP4 interaction. The second series of AKR-XI-85 analogs included compounds in which the 2-trifluoromethylphenyl head group was replaced (Table 4). Despite the favorable docking scores, heterocyclic head groups (**98-103**) completely eliminated all *in vitro* activity. The compound with an unsubstituted phenyl head group (**85**) and all derivatives with a 3-substituted phenyl ring (**81-84, 86**) showed significantly decreased or no *in vitro* potency. It is interesting to note that while the presence of the 2-trifluoromethyl group seems critical for RBP4 binding, the 3-trifluoromethylphenyl analog (**86**) displayed no activity. The replacement of the trifluoromethyl group in the C-2 position with substituents of varying sizes and polarities (**87-92**) had a detrimental effect on the potency with a few notable exceptions, the 2-trifluoromethoxy (**93**), 2-pentafluoroethyl (**95**), and 2-isopropyl (**94**) groups. Substitution with polar groups was particularly poorly tolerated, regardless of the substituent’s size.

**Fig. 6.**
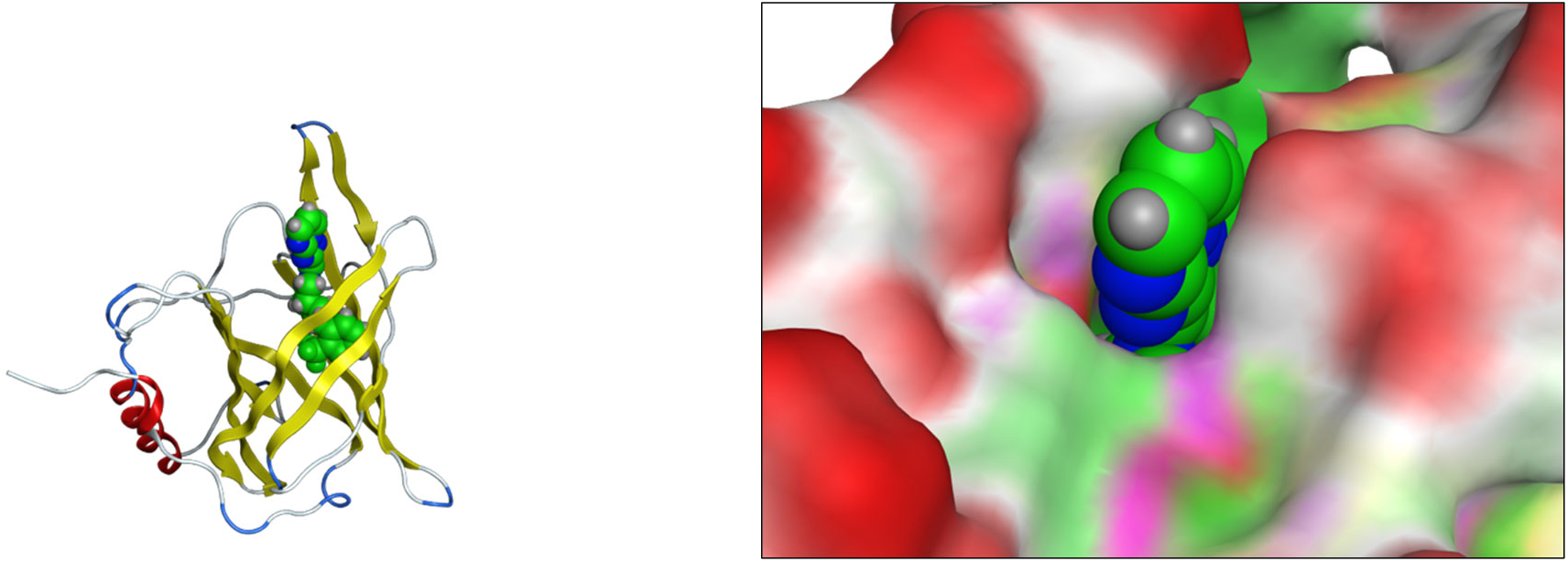
Docked conformation of AKR-XI-85 (**40**) in the binding pocket of RBP4 and key receptor interactions. The left panel shows the situation of AKR-XI-85 in the RBP4 interior, and the right panel shows the portion that is exposed to solvent. For clarity, AKR-XI-85 is shown in space-filling representation. RBP4 structure taken from PDB entry 3FMZ.

Prior to the *in vivo* testing of AKR-XI-85, we conducted an *in vitro* metabolic stability analysis in four species (mouse, rat, non-human primate, and dog) as well as an additional *in vitro* ADMET evaluation (Tables 5 and 6). The compound displayed reasonable stability in the mouse and cyno microsomal stability assays and good-to-excellent stability in the dog and human microsomal assays. AKR-XI-85 exhibited reduced metabolic stability in rat liver microsomes, suggesting a potential for lower systemic exposure in rats after systemic administration. This could pose a challenge for utilizing rats as the preferred rodent species in GLP-compliant safety assessment studies. The metabolite identification study conducted in human, rat, mouse, dog, and cynomolgus monkey liver microsomes revealed no human-specific metabolites. This suggests that safety assessment in any of the common safety species would be adequate for predicting human toxicities.

AKR-XI-85 lacked limiting inhibitory activity in a standard CYP 450 panel, showed no hERG inhibition and no PXR activation (Table 6), demonstrating overall good drug-like characteristics.

AKR-XI-85 testing in the broad Cerep-Panlabs selectivity panel revealed no obvious off-target activities. Permeability assessments conducted in MDCKII-MDR1 and Caco-2 cells suggest that AKR-XI-85 is a highly permeable compound, indicating its anticipated strong *in vivo* absorption. The compound does not seem to be a substrate for the P-glycoprotein efflux transporter.

The pharmacokinetic studies conducted in mice and dogs demonstrated that in mice AKR-XI-85 displayed reasonably high oral bioavailability but rapid clearance after oral and intravenous administration (Table 7). Because of rapid systemic clearance, modest AKR-XI-85 exposure was achieved in mice. Consistent with the results of the dog microsomal stability assay, AKR-XI-85 displayed low clearance and a high exposure in dogs, demonstrating excellent PK characteristics in this species. Given the modest exposure achieved in mice, it was crucial to assess the pharmacodynamic effects following the oral administration of AKR-XI-85 in this species. Serum RBP4 lowering in response to compound administration provides a convenient PD marker for evaluating the target engagement and *in vivo* potency of RBP4 antagonists, such as AKR-XI-85.

We administered AKR-XI-85 orally to mice at increasing concentrations (15, 25, or 35 mg kg^−1^) to test the ability of the compound to lower the levels of serum RBP4. We observed a robust dose-dependent reduction in levels of circulating RBP4 at all three doses, with the highest oral dose resulting in a maximum reduction of 89% (Fig. 4). The RBP4-lowering effect was durable with a 25% decrease still observed at the 24-hour time point after dosing at the 35 mg kg^−1^ dose. These pharmacodynamic results align well with the impressive *in vitro* potency of AKR-XI-85 against the target. They also confirm that the compound exposure attained in mice is sufficient to elicit the desired magnitude of the PD response. The effect of AKR-XI-85 on the accumulation of toxic bisretinoids was next studied in the *Abca4^-/-^*mouse model of Stargardt disease. This model is recognized as an established standard for evaluating the preclinical efficacy of bisretinoid-lowering therapies. The genetic ablation of *Abca4^-/-^* renders these animals incapable of transporting *N*-retinylidene-PE across the disk membranes efficiently, leading to an elevated accumulation of lipofuscin bisretinoids. (61) AKR-XI-85 performed well in the *Abca4^-/-^* mice over the 60-day dosing period inducing a sustained 72% serum RBP4 reduction (Fig. 5A). This RBP4 lowering induced by AKR-XI-85 resulted in a robust 70% reduction in the A2E concentration in the retinas of *Abca4^-/-^*mice compared to the vehicle-treated animals with the same genetic background (Fig. 5B).

In summary, our study demonstrated that the RBP4 antagonist triazolopyrimidine AKR-XI-85, which lacks undesired off-target PPARγ agonistic activity, effectively induces a significant reduction of circulating RBP4 in mice. The chronic administration of AKR-XI-85 in the mouse model of Stargardt disease resulted in a sustained reduction of serum RBP4 and a substantial 70% reduction in measured lipofuscin bisretinoids in the eyes of treated animals. The favorable *in vitro* and *in vivo* potency of AKR-XI-85 along with its appropriate PK characteristics justify further development of this compound and indicate that AKR-XI-85 holds potential as a therapeutic option for treating Stargardt disease and related lipofuscin-dependent retinal dystrophies.

## Acknowledgments

This study was supported by a Gund-Harrington National Initiative Award TA-NMT-1115-0690-COLU-GH to K.P.

S.-X.D., A.R., A.S.W., and D.W.L. would like to thank the NIH for providing funds for their LC/MS (1S10OD018121) and the DoD for providing funds for their NMR spectrometer (N00014-11-1-0900).

